# Accurate and simultaneous identification of differential expression and splicing using hierarchical Bayesian analysis

**DOI:** 10.1101/2019.12.16.878884

**Authors:** Guy Karlebach, Peter Hansen, Diogo F.T. Veiga, Robin Steinhaus, Daniel Danis, Sheng Li, Olga Anczukow, Peter N. Robinson

## Abstract

The regulation of mRNA controls both overall gene expression as well as the distribution of mRNA isoforms encoded by the gene. Current algorithmic approaches focus on characterization of significant differential expression or alternative splicing events or isoform distribution without integrating both events. Here, we present Hierarchical Bayesian Analysis of Differential Expression and ALternative SPlicing (HBA-DEALS), which simultaneously characterizes differential expression and splicing in cohorts. HBA-DEALS attains state of the art or better performance for both expression and splicing, and allows genes to be characterized as having differential gene expression (DGE), differential alternative splicing (DAST), both, or neither. Based on an analysis of Genotype-Tissue Expression (GTEx) data we demonstrate the existence of sets of genes that show predominant DGE or DAST across a comparison of 20 tissue types, and show that these sets have pervasive differences with respect to gene structure, function, membership in protein complexes, and promoter architecture.

## INTRODUCTION

RNA sequencing (RNA-seq) has become the most commonly used genomic technique for the transcriptome-wide analysis of differential expression and alternative splicing of mRNAs. Since its introduction over a decade ago, Illumina short-read sequencing technology has been the dominant platform for carrying out RNA-seq experiments, but newer long-read single-molecule sequencing technologies of Pacific Biosciences and Oxford Nanopore provide alternatives that may allow a more accurate and comprehensive assessment of isoform diversity. ^1^ Analysis of RNA-seq data is done in a pipeline that maps raw reads to genes or isoforms (transcripts), quantifies the number (count) of reads associated with each isoform generating an expression matrix, followed by normalization steps and statistical analysis of differential expression. ^2;3^

Algorithms for the analysis of differential gene expression or differential splicing have many different approaches. Differential gene expression (DGE) refers to alterations in the expression (counts) of the sum of each of the isoforms that are encoded by a gene. Many methods for DGE analysis are based on discrete probability distributions such as the Poisson or negative binomial. ^4;5^ voom instead estimates the mean-variance relation non-parametrically from log-counts per million reads data, which are used as input for linear modeling and empirical Bayes differential expression analysis. ^6^

In contrast to DGE, differential alternative splicing and transcription (DAST) refers to differential usage of isoforms that include distinct combinations of exons or begin from distinct transcription start sites. Computational methods for identifying DAST in RNA-Seq data can be broadly divided into two approaches. The first approach is based on an analysis of the percent spliced in (Ψ[Psi]), which is defined as 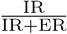, where IR refers to inclusion reads and ER to exclusion reads. This approach models differences in splicing as differences in Ψ, corresponding to the probability of an alternative splicing event at a splice junction. ^7–10^ A second approach compares counts of alternative isoforms. ^11–13^

Most existing methods look at either DGE or DAST but not both. When using separate procedures for differential splicing and expression, however, the determination of which genes are alternatively spliced, differentially expressed, undergo both of these changes or none of them requires intersection between negative and positive findings. This is usually not possible without violating some of the assumptions of individual tests. For example, frequentist methods assess statistically significant differential expression or splicing using *p*-values, but non-significant *p*-values cannot be readily interpreted as providing evidence of lack of differential expression or splicing. Moreover, variation of gene expression can be affected by expression level, ^14^ and as a corollary variation of isoform expression is affected by isoform level. Methods that transform isoform levels into proportions (i.e., Ψ-based methods) do not model this relationship, and therefore fail to accurately model dispersion, which is essential for determining significance. On the other hand, modeling individual isoform expression levels but not modeling their joint expression can result in false positives when a gene is differentially expressed but its isoforms are not differentially spliced. Similarly, a change in an individual isoform’s levels between conditions does not necessarily mean a change in that isoform’s proportion and vice versa.

In this work, in contrast, we present a Bayesian method for analyzing RNA-seq data that simultaneously identifies DGE and DAST based on isoform counts. We show with our method, using data from the Genotype-Tissue Expression (GTEx) project, ^15^ that genes can be assigned to four groups according to the propensity of a gene to show DGE, DAST, both, or neither in comparisons between different tissue types. These classes differ not only with respect to gene functions and structure, but also with respect to the distribution of transcription factor binding sites (TFBS), membership in protein complexes, and methylation of their promoter regions.

### RESULTS

Here, we present a method for joint modeling of differential expression and splicing, Hierarchical Bayesian Analysis of Differential Expression and ALternative Splicing (HBA-DEALS).

### A Bayesian model for simultaneous assessment of differential expression and splicing

HBA-DEALS uses as input a matrix of isoform counts derived from mapped RNA-seq reads in *n* samples divided into two cohorts *n*_1_ and *n*_2_ (e.g., cases and controls). HBA-DEALS uses a hierarchical Bayesian model that simultaneously assesses the absolute expression levels of the gene and its isoforms (Fig. 1).

**Figure 1.**
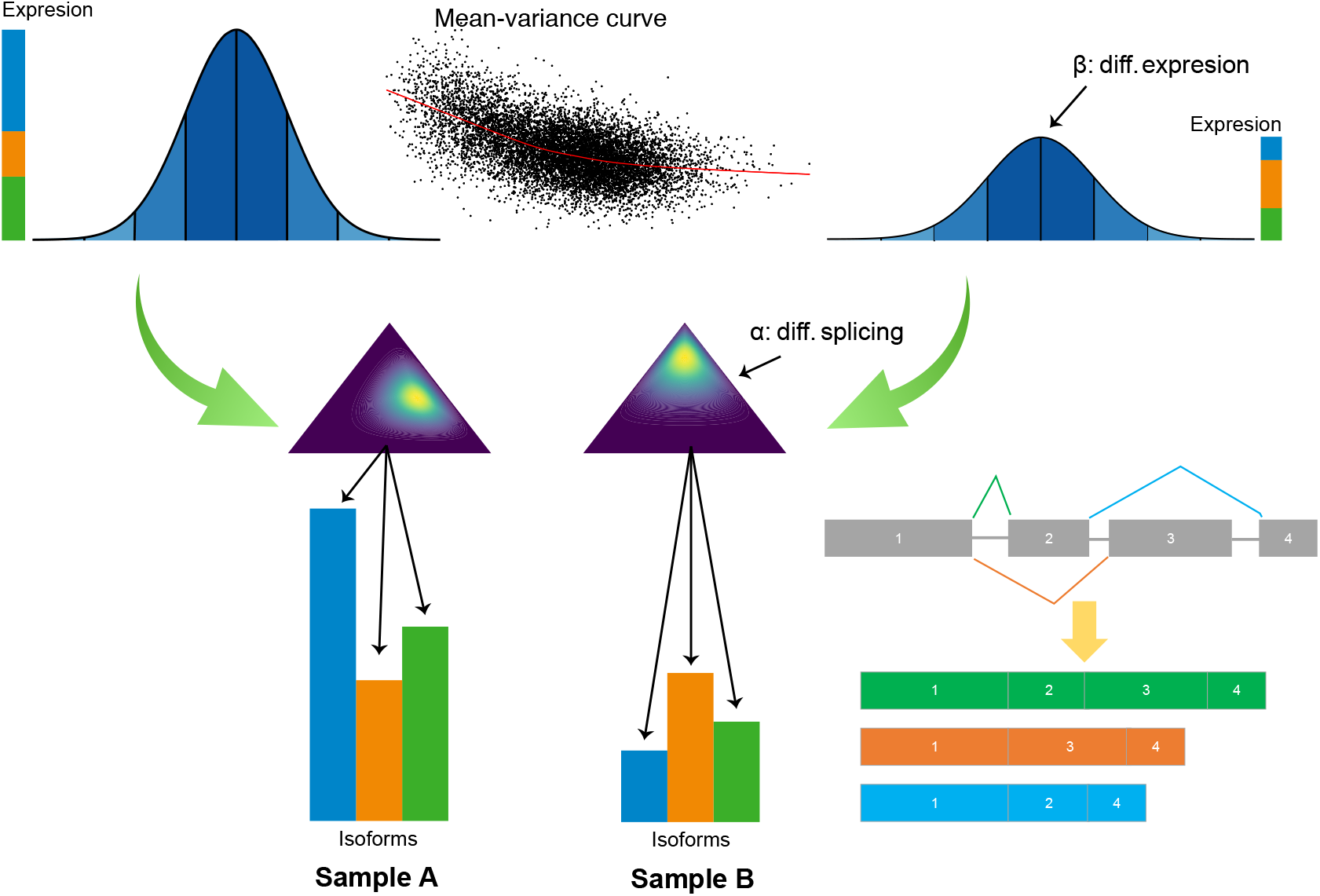
Overview of the HBA-DEALS model. The expression of a gene with three isoforms (green, orange and blue) is shown. The gene expression is the sum of the expression of the isoforms. Differential gene expression is modeled as two Normal distributions that differ by the parameter *β*. The expressions of the corresponding isoforms are modeled using a Dirichlet prior whose difference is controlled by *α* (symbolized by the two triangles). An MCMC procedure is used to solve for the posterior distribution of the parameters of the model for all genes and isoforms at once. The posterior distributions are in turn interpreted to classify each gene as DAST, DGE, DAST/DGE, or static (Methods).

The isoform counts are first log-transformed. Gene expression levels are then modeled using a normal distribution, with mean that is equal to the sum of corresponding mean isoform levels. A linear model is fit to each gene’s levels, and a trend line is then fitted to the square root standard deviations as a function of mean gene level. ^6^ The variance of an individual sample is inferred from the value of the corresponding value of fitted function. The mean isoform expression is a fraction of the mean expression level of its gene (In Fig. 1, Sample A and Sample B display a different distribution of the three isoforms of an example gene), and similarly to gene expression, a sample-specific variance is obtained from a mean-variance trend for isoforms. The proportions of all isoforms assigned to a gene are represented as a vector of isoform fractions [*p*_1_*, p*_2_, .., *p_n_*] with ∑ *p_i_* = 1 (isoform fractions are symbolized by the triangles in Fig. 1).The prior of the isoform fractions is Dirichlet distributed with the vector [1, 1, …, 1].

In order to model difference in gene expression, a parameter *β* is added to the mean expression level in one condition. A non-informative 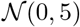 prior is assigned to *β*. The model predicts DGE when the bulk of the posterior of *β* does not contain 0. Unlike differential expression, simple addition cannot model difference in splicing because the entries of the vector of isoform fractions must sum to 1, and addition does not preserve this property. Therefore, instead of adding a vector *α* to the isoform fractions in one condition, we apply an Aitchison perturbation ^16^ between *α* and the isoform fractions in that condition. As with *β*, HBA-DEALS estimates the posterior distribution of *α*, and uses this to call DAST (Methods). Given the expression levels of a gene, the isoform proportions are independent of *β*. Thus, HBA-DEALS independently and simultaneously assesses DGE and DAST.

### Model validation

We applied three approaches to assess the performance of HBA-DEALS. First, we extended an existing simulation scheme for RNA-seq expression data ^6^ to enable the modeling of alternative splicing. For each gene, we split its sample proportion between a random number of isoforms, and for differentially spliced genes we doubled the proportion of one random isoform in cases and another in controls. We analyzed 50 simulated datasets for DGE with HBA-DEALS, voom, ^6^ DESeq2, ^5^ edgeR^4^, baySeq ^17^, and NOISeq. ^18^ We analyzed the same datasets for differential alternative splicing with HBA-DEALS, rMATS, ^8^ and a method we call optimal splicing from transcript proportions (OSTP), which provides an upper bound on the performance of t-statistic-derived significance values to compare Ψ of isoforms (Methods). HBA-DEALS displayed a larger area under the precision-recall curve than the other approaches did for both DGE and DAST (Fig. 2a-b).

**Figure 2.**
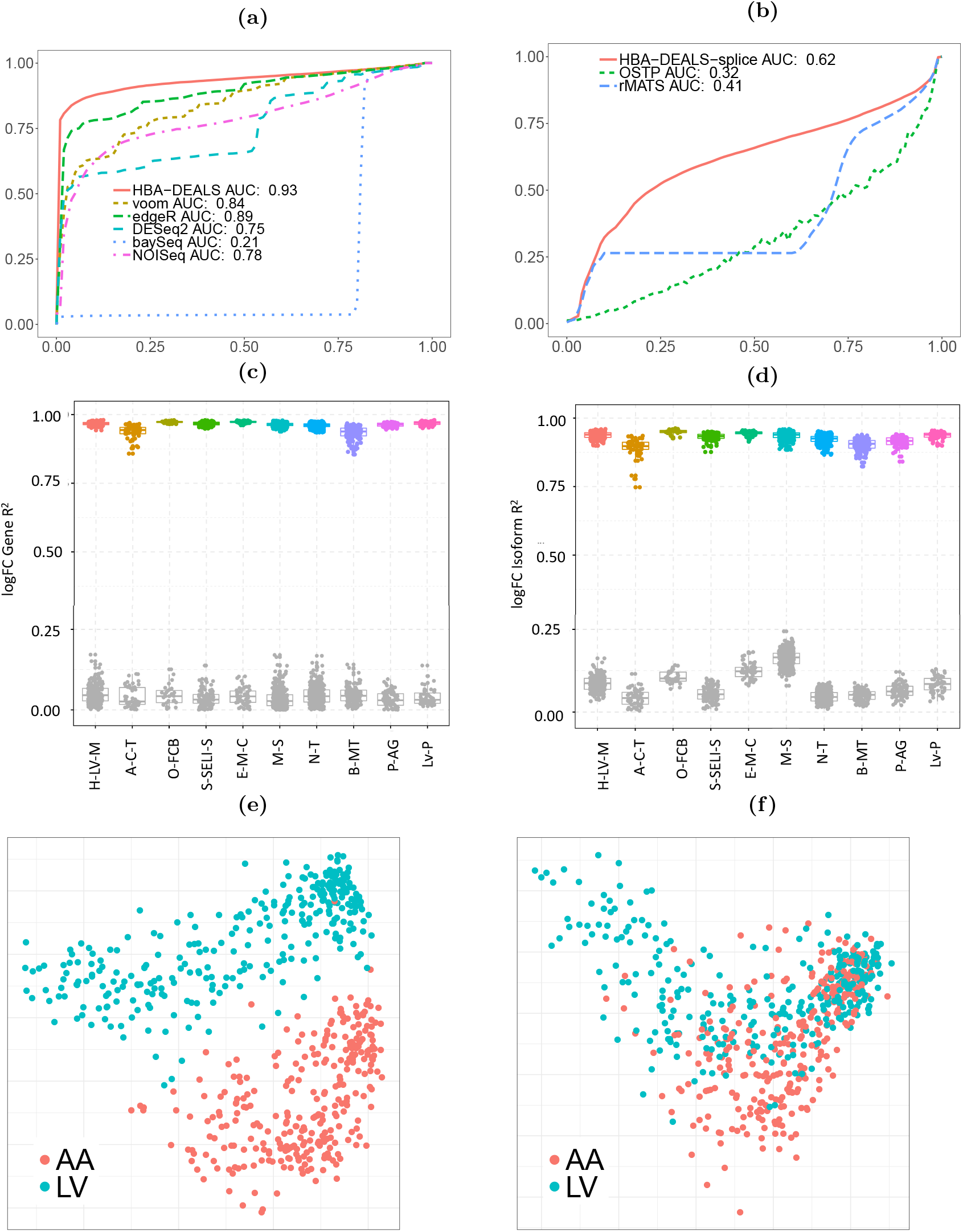
HBA-DEALS evaluation. **(a)** Precision-Recall (PR) analysis of HBA-DEALS and five state-of-the-art algorithms for the detection of RNA-seq differential expression. **(b)** PR analysis of HBA-DEALS, OSTP, and rMATS. **(c)** *R*^2^ for gene expression (GTEx, distant). Data points shown in color represent correlations based on genes that were identified as differentially expressed by HBA-DEALS. As a control, genes not identified as differentially expressed (probability of direction *p* ≥ 0.25, see Methods) are shown in gray (Abbreviations in Supplemental Table S2). **(d)** *R*^2^ for isoform proportion (GTEx, distant). Data points shown in color represent correlations based on isoforms identified as differentially spliced by HBA-DEALS. As a control, isoforms from genes not identified as differentially spliced (*p* ≥ 0.25) are shown in gray. **(e)** MDS plot for left ventricle (LV) compared with atrial appendage (AA). Each dot represents one sample. The MDS was performed using only isoforms that HBA-DEALS identified as DAST. **(f)** As a control, the same analysis was performed using isoforms of genes from the DGE group.

In order to assess the accuracy of HBA-DEALS using real data, we ran HBA-DEALS on estimated isoform levels from the Genotype-Tissue Expression project (GTEx)^15^. We used HBA-DEALS to identify differentially expressed and differentially spliced genes in 20 different pairs of tissues. We chose ten pairs of tissues that were closely related (e.g., subcutaneous adipose tissue and visceral adipose tissue), and ten that were more distant (e.g., liver and pituitary gland; Supplemental Table S1). We formed multiple subcohorts for each tissue by choosing 15 samples at random and then compared the results of HBA-DEALS between different subcohorts. Although the individual samples derive from unrelated individuals, there was a high degree of overlap of genes identified as differentially expressed or spliced. We tested the overlap between a total of 2791 pairs of cohorts (Supplemental Table S1). In each case, the overlap was highly significant (*p* < 2.23 × 10^−308^ for all comparisons, hypergeometric test). These results suggest that HBA-DEALS is able to identify characteristic and reproducible differences in cohorts (Fig. 2c-d and Supplemental Figs. S1 and S2). The correlation was higher in the distant tissues, likely because ratio of larger than the differences between samples was higher. We also verified that the correlation increases with cohort size (Supplementary Fig. S3).

Finally, for each tissue pair we performed multidimensional scaling (MDS) of the vectors of isoform proportions in all available samples, including only isoforms that were differentially spliced in at least 3 subcohorts (Methods). More specifically, we first obtained genes that were predicted to be differentially spliced in at least 3 cohorts of a given tissue pair, and their isoforms that were classified as differentially spliced with a fold change of at least 2. For each gene we then computed the proportion of each isoform that was predicted to be differentially spliced out of the total number of isoforms of that gene. The Euclidean distance between the vectors of these proportions over all differentially spliced genes of a tissue pair were then reduced into 2 dimensions using multidimensional scaling. ^19^ The overlaps were highly significant (*p* < 2.23 × 10^−308^, Hypergeometric test). As an example, we show MDS that was performed on samples from the left ventricle of the heart (LV) and the atrial appendage (AA). We first performed MDS by calculating intersample distance based on levels of isoforms that had been identified as differentially spliced in different cohorts and obtained a nearly perfect separation by tissues (Fig. 2e). As a control, we repeated the MDS with differentially expressed genes and isoforms that were assigned probability of at least 0.25, and the two cohorts display a substantially lower degree of separation (Fig. 2f).

### HBA-DEALS defines four categories of genes that differ with respect to splicing and expression

We next asked whether sets of genes can be identified by HBA-DEALS whose regulation is found to occur primarily by means of differential splicing, differential expression, or both. For the following analysis, we performed 20 comparisons between samples from different tissues (e.g., left ventricle against atrial appendage). For each comparison, we compared cohorts of 15 samples for each tissue (Supplemental Table S1), and used HBA-DEALS to call genes differentially spliced, differentially expressed, both, or neither.

For the following analysis, a gene is considered differentially expressed in a tissue if it was differential expressed in at least 3 subcohorts, and differentially spliced if it has an isoform that was differentially spliced in at least 3 subcohorts. If a gene is found to be differentially spliced in at least twice as many comparisons as it is found to be differentially expressed, we assign it to the DAST group. Conversely, if a gene is differentially expressed in at least twice as many comparisons as it is differentially spliced, we assign it to the DGE group. Genes that are both differentially spliced and differentially expressed are assigned to the DAST/DGE group. Finally, genes that do not display differential expression or splicing as defined above are assigned to the static group (Fig. 3a).

**Figure 3.**
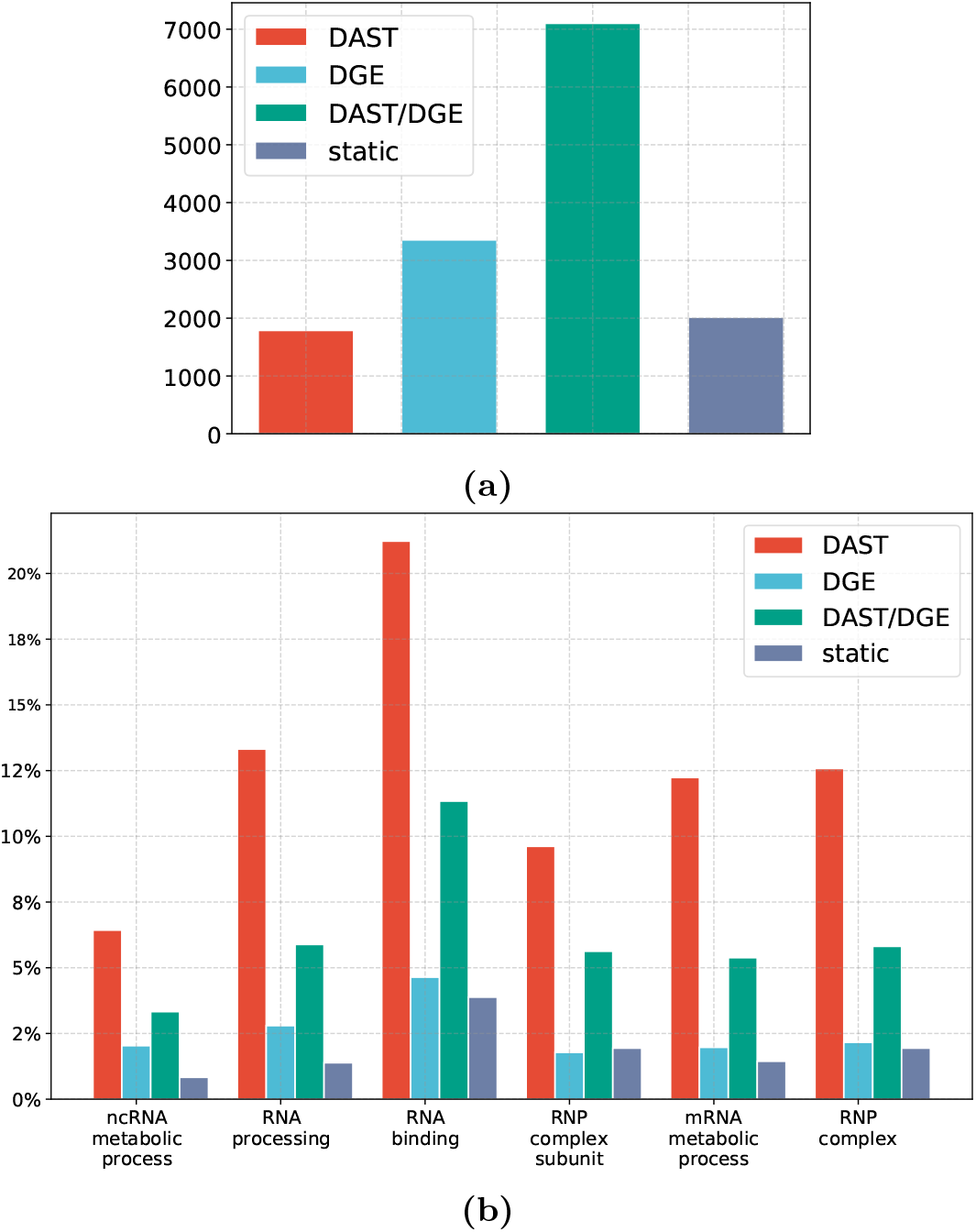
**(a)** Distribution of protein coding genes in the groups DAST (n=1789), DGE (n=3355), DAST/DGE (n=7102), and static (n=2018). Only those genes were included in the analysis for which at least two isoforms were expressed at sufficient depth in most tissues. **(b)** Gene Ontology analysis revealed six GO significantly enriched GO terms that show an at least 2-fold enrichment in one of the four groups (Methods and Supplemental Tables S3–S8).

We performed Gene Ontology ^20^ (GO) term enrichment analysis on the genes in each of the four groups. The four groups differed with respect to significantly overrepresented GO terms (Fig. 3b, with details in Supplemental Tables S3–S8). Six terms displayed strong enrichment in one of the four classes, as defined by statistically significant overrepresentation, having at least 20 annotated genes, and showing an at least two-fold higher percentage of annotated genes than the entire population of genes (Methods). All six enrichments were found for the DAST group, and all of the terms were related to RNA biology. For instance, while only 10.1% of all 13,688 genes were annotated to rna binding, over twice as many genes in the DAST set were (21.5%). RNA binding proteins are involved in each step of RNA metabolism including alternative splicing. ^21^ Changes in alternative splicing are common in biological processes and disease states, and an investigation of the functions of alternatively spliced genes may tell us something about the biology of those states. For example, subsets of alternatively spliced genes found in aging, with mutations in the spliceosome gene *U2AF1* in myelodysplastic syndrome, and with differentiation of erythroblasts are enriched for genes involved in RNA processing. ^22–24^

None of the significant GO terms for the DGE group was associated with a two-fold increase in the percentage of annotated genes. The specificity of the significant GO terms identified for the DAST and DGE groups was significantly higher than for the remaining two groups (Supplemental Fig. S4). This suggested that the DAST and DGE groups show a greater degree of functional uniformity than the other two groups, which motivated us to further investigate differences between these two groups.

### Pervasive differences in the genomic characteristics of DAST and DGE genes

We compared the DAST and DGE groups with respect to a variety of genomic properties. DAST genes show a lower percentage of promoter methylation than DGE genes. DAST genes also have a higher number of exons, are shorter and have a lower mean exon length. DGE genes are more likely to have a TATA-box element in their promoter, and are correspondingly less likely to be associated with a CpG island. DAST and DGE genes differ with respect to the frequency of a number of predicted transcription factor binding sites (Table 1).

**Table 1.** Genes regulated predominantly by alternative splicing (AS genes) differ from genes regulated predominantly by differential expression (DE genes) with respect to a number of characteristics related to DNA sequence, methylation, and transcription factor binding. Entries shown as percentages indicate the percentage of all genes in the group that display the characteristic. MW: Mann-Whitney test; FET: Fisher exact test; W: Wald test. Additional methylation results are listed in Supplementary Table S9.

| Predictor | DAST genes | DGE genes | *P*-value |
| --- | --- | --- | --- |
| Promoter methylation age 0 | 34.66 | 25.09 | $1.69 \times 10^{-23}$ (MW) |
| ZFX binding | 57.5% | 35.8% | $1.83 \times 10^{-8}$ (W) |
| Number of Exons | 34 | 21 | $4.22 \times 10^{-106}$ (MW) |
| Total Gene Length | 72,367 bp | 76,408 bp | $5.81 \times 10^{-6}$ (MW) |
| Dispersion | 27.32 | 27.92 | $6.62 \times 10^{-3}$ (MW) |
| CpG island | 77% | 61.3% | $1.52 \times 10^{-26}$ (FET) |
| TATA box | 5.5 % | 12.9% | $2.32 \times 10^{-15}$ (FET) |
| Number of transcripts | 11.4 | 6.9 | $4.94 \times 10^{-152}$ (MW) |
| Exon length | 240 bp | 250 bp | $4.35 \times 10^{-28}$ (MW) |

DNA methylation of the promoter region or of the gene body can influence alternative splicing. ^25–27^ We compared the methylation levels in gene body and promoter of DAST and DGE genes, in 12 different age groups (Methods). In all the age groups, the percentage of methylation was significantly higher in expression-regulated genes. We found that the degree of promoter and gene body methylation is also correlated with exon count of genes. The proportion of DAST genes increases with the exon count, but for any particular exon count, lower degrees of promoter methylation are association with higher proportions of DAST genes (Fig. 4a). For gene body methylation, the proportion of DAST genes is highest with intermediate levels of methylation, and lowest with the lowest levels of methylation. Furthermore, when the number of exons grows, even small increases in methylation increase the chance that a gene is splicing-regulated (Fig. 4b).

**Figure 4.**
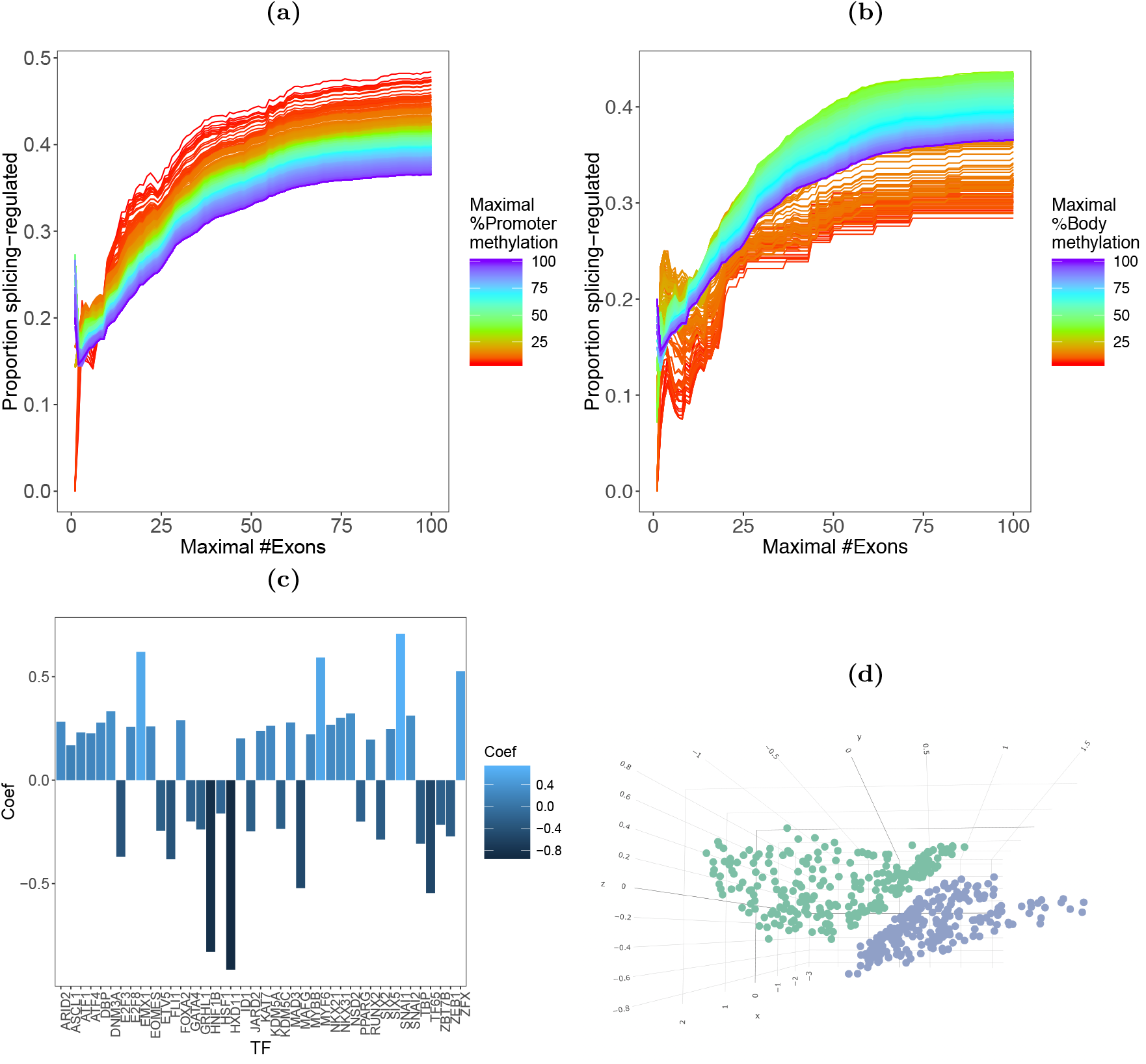
**(a)** Promoter Methylation × Number of Exons × Proportion of Splicing Genes - Promoter. **(b)** Promoter Methylation × Number of Exons × Proportion of Splicing Genes-Gene Body. **(c)** Bar plot of the 41 TFs with significant regression coefficients in the logistic regresssion. Positive coefficients predict DAST, and negative coefficients predict DGE. **(d)** Multidimensional Scaling of TF interaction profiles of DAST (blue) and DGE (green) genes.

Limited evidence exists coupling binding of transcription factors to promoters with alternative splicing. ^28;29^ We explored whether profiles of TF binding motifs in gene promoters are predictive of a gene being in the DAST or DGE group by means of logistic regression with a total of 401 predictors consisting of the predicted target genes of 401 TFs. The value of each predictor is 1 (TFBS present) or 0 (no TFBS). The dependent variable is the group (DAST vs. DGE). The weighted sum of the predictors and the intercept models the logit of the probability of belonging to the expression-regulated gene class. The model identified 41 TFs with a statistically significant regression coefficients, comprising 24 TFs in the DAST group and 17 TFs in the DGE group (Fig. 4c and Supplemental Table S10). We then used the Mann-Whitney test to compare the distributions of the probability of a gene belonging to the DAST class as assigned by the network to the genes in the DAST and DGE classes. We obtained p-values of 1.11 × 10^−76^ and 5.25 × 10^−77^. This indicates that the model can predict the correct gene class for unobserved genes based in TF binding and methylation profiles.

### Networks of splicing-regulated transcription factors

We then investigated potential synergy between TFs of the DAST and DGE groups. For each group and for each pair of TFs, we computed the number of targets bound by both TFs divided by the number of targets bound by at least one of them, where targets are defined as all genes in either the DAST or the DGE groups. We noted that some TFs had very few or very many binding targets. In order to remove trivial low and high scores we therefore selected TFs from the 0.2 to the 0.8 quantiles with respect to the total number of targets in both sets of genes. We then performed multidimensional scaling on the vectors of interaction scores of the different TFs, i.e. each point in the MDS corresponds to the interaction profile of a specific TF with other TFs when the targets are either the DAST or DGE set (Fig. 4d). Remarkably, we obtained perfect linear separation between interaction profiles for the two types of genes. This result suggests that combinatorial regulation plays a role in determining both changes in splicing and gene expression.

In order to examine whether particular protein complexes are enriched for DAST, we downloaded the full set of gene complexes from the CORUM database ^30^ and computed the probability of the obtaining the observed proportion of DAST genes or a higher proportion under the binomial null distribution, setting the probability of a DAST gene to the mean proportion of DAST genes over all complexes (0.55). After Benjamini-Hochberg multiple testing correction, DAST but not DGE were significantly enriched in spliceosome related complexes. Two complexes enriched for DAST genes were related to the ribosome (Table 2). Interestingly, networks of autoregulated alternative pre-mRNA splicing have been demonstrated for components of the spliceosome as well as for subsets of ribosomal proteins. ^31–33^

**Table 2.**
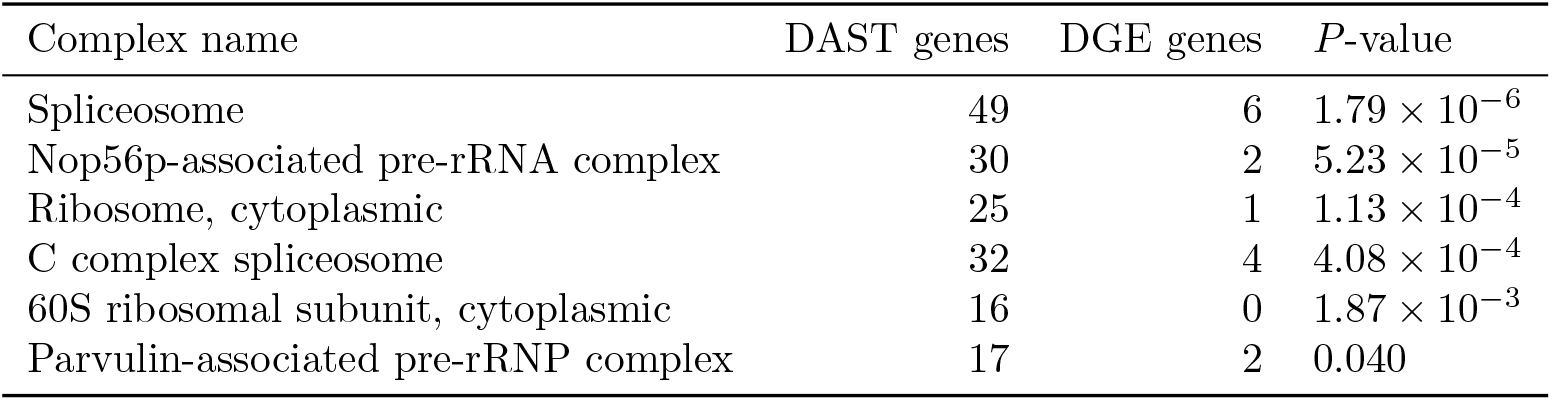
Enrichment of DAST genes in protein complexes was calculated using the binomial distribution (Methods). The Benjamini-Hochberg-corrected *p*-value is shown.

Combinatorial interactions among transcription factors (TFs) and TF subnetworks are critical for tissue-specific gene expression. ^34^ We therefore asked to what extent transcription factors represented in the DAST and DGE groups differ. We computed the count of DAST-TFs with TF binding motifs in promoters of all DAST genes and the corresponding count with motifs in promoters or DGE genes. There was a significantly higher count for DAST genes as compared to DGE genes (16.05 vs 13.67 binding TFs, *p* = 7.36 × 10^−53^, Mann-Whitney test). This finding supports the possible existence of independent cellular circuits that are based primarily on changes in alternative splicing (Supplemental Fig. S5).

## DISCUSSION

Although short- and long-read RNA sequencing has become a standard method for characterizing both gene expression and alternative splicing, the interplay between gene expression and alternative splicing has not been extensively studied. The majority of existing methods interrogate either gene expression or alternative splicing but not both, and methods for jointly modeling expression and splicing have been lacking. In this work we have presented HBA-DEALS, a Bayesian method that analyzes both splicing and expression in a single model. Using simulated data we have shown that the performance of HBA-DEALS in the identification of differential gene expression and differential alternative splicing is superior to that of state-of-the-art approaches. By investigating data of the GTEx project, we additionally showed that the predictions of HBA-DEALS are reproducible across independent biological cohorts. The algorithmic approach HBA-DEALS is a paradigm that can be used to identify groups of genes that display DAST, DGE, both, or neither. This analysis result is not readily available with existing methods that investigate DGE or DAST one at a time.

To demonstrate a typical application of our algorithm, we characterized a set of genes that are preferentially alternatively spliced over a large number of comparisons of different tissues using data from the GTEx project. We characterized a set of genes that are preferentially alternatively spliced across comparisons of 20 tissue types. The DAST set was enriched in functions related to RNA metabolism, for members of spliceosomal and ribosomal protein complexes, and showed pervasive differences compared to the DGE set with respect to gene structure (gene length, average exon length, total exon count), DNA methylation of both promoter and gene body sequences, as well as the distribution of transcription factor binding sites. We have further found that combinatorial regulation of genes by transcription factors is fundamentally different in splicing-regulated and expression-regulated genes, which suggests that both processes are under the control of different gene regulatory networks. It seems plausible that each process can act independently under certain conditions or given a specific set of triggers. In support of this hypothesis, we found that transcription factors that are themselves in the DAST group favor targets that are in the DAST group. Moreover, DAST genes are enriched for RNA-binding, suggesting possible post-transcriptional regulatory interactions.

HBA-DEALS is an R package that is freely available at https://github.com/TheJacksonLaboratory/HBA-DEALS.

## METHODS

### HBA-DEALS: Hierarchical Bayesian Analysis of Differential Expression and ALternative Splicing

Hierarchical Bayesian modeling (HBM) is a multiparameter modeling technique in which one assumes a statistical distribution for individual parameters whose interdependencies are reflected in the structure of the hierarchy. In HBA-DEALS, this hierarchy models isoform levels as fractions of the total number of mRNA molecules produced from a certain gene. Our assumptions in developing our model were: (i) Increase or decrease in gene expression induces increase or decrease in the level of at least one isoform; (ii) Isoform levels are fractions of the gene expression level; and (iii) Changes in isoform fractions do not necessitate changes in expression levels and vice versa. A Markov Chain Monte-Carlo (MCMC) technique can be used to estimate the posterior probability of the parameters of an HBM. To do so, one must design the structure of the HBM and define the probability distribution of each node.

The input for HBA-DEALS consists of a matrix of gene and isoform counts derived from to two different conditions, here referred to as *case* and *control*, using any short- or long-read next-generation sequencing technology. In the case of short-read RNA-seq, tools such as RSEM ^35^ or StringTie, ^36^ can be used to calculate isoform counts. Long-read isoform counts can be generated with pipelines such as SQANTI. ^37^ HBA-DEALS calculates gene expression levels by summing up the isoform counts of individual genes.

The data is log-transformed using *log*_2_(*x* + 0.5), where *x* is the count-per-million reads (log-cpm). Expression levels are modeled as Normal, with mean that is the sum of the corresponding isoform levels, and sample-specific variance that is obtained from a mean-variance trend by fitting a linear model to each gene’s levels, and then fitting a trend line to the square root standard deviations as a function of mean gene level. ^6^

### Assessment of DGE

HBA-DEALS models gene expression as follows. The difference between cases and controls is modeled with a non-informative Normal prior.

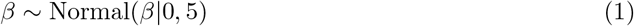

The mean log-cpm level in cases is modeled with a Normal distribution such that the mean of its prior is equal to the mean expression value of the control samples (*μ*_1_) with a variance of 5.

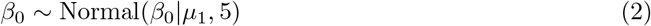

In order to model differences between cases and controls, we model the expression in control *i* (*y_i_*) as:

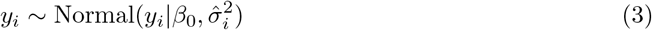

For case *j*, the mean is defined as *β*_0_ + *β*:

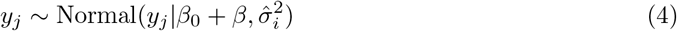

### Probability of direction

If the mean of the posterior is greater than 0, the probability of differential splicing is 1 minus the posterior density that is smaller than 0. If the mean is smaller than 0 the probability is 1 minus the posterior density that is greater than 0. This is essentially the probability of direction with mean in place of the median of the posterior. The mean of the intercept for expression levels is the log-transformed mean in the control group. For MCMC initialization, gene expression values are set to the log-transformed sum of isoform counts and *β* is set to 0. *β* is set to 0.

### Assessment of DAST

Isoform levels are also modeled as normal, with variance that is obtained from a mean-variance trend similarly to expression levels. The mean of isoform *i* corresponds to a fraction *p_i_* of the mean expression level, with ∑ *p_i_* = 1.

The control fractions have a *Dirichlet*(1) prior, and the case fractions relate to the control fractions (*p_i_* and 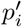 refer to the proportion of isoform *i* in controls and cases) via the following formula (the Aitchison perturbation ^16^):

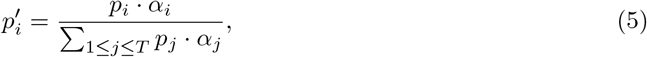

where *T* is the number of isoforms of the gene and *α* is a vector whose entries sum to 1.

This Aitchison perturbation computes the product of corresponding entries in the two vectors, and divides each entry in the resulting vector by the sum of its entries. For example, if a gene has two isoforms with fractions (0.4,0.6) in condition 1, and alpha is (0.6,0.4), then the Aitchison perturbation will map the fraction of each isoforms to 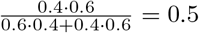 in condition 2. A vector whose entries are equal and sum to 1 is the identity element of the Abelian group defined on the simplex by the Aitchison perturbation. Therefore, a gene is not differentially spliced if and only if all the entries of *α* are equal. The prior on *α* is also set to Dirichlet with the vector 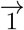.

For MCMC initialization, we set the initial values of *p_i_* to

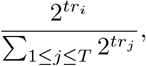

where *tr_i_* is the log-cpm level of the *i_th_* isoform. Additionally, *α*’s entries are initialized to 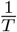.

If for the *i^th^* isoform most of the posterior density of *α_i_* is concentrated away from the scalar product ***α* · p**, then that isoform is predicted to be differentially spliced. If the posterior density of any isoform of a gene is concentrated away 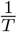, the gene is predicted to be differentially spliced. Therefore, we predict differential splicing of a specific isoform when the bulk of the marginal posterior density of the isoform’s corresponding entry in *α* does not contain the scalar product of *α* and the vector of isoform proportion in condition 1. We predict differential splicing of a gene when the bulk of the marginal posterior density of any of its isoforms does not contain 1 divided by the number of isoforms. More specifically, if the mean of the posterior is greater than the threshold value the probability of differential splicing is 1 minus the posterior density that is smaller than the threshold, and if the mean is smaller the probability is 1 minus the posterior density that is greater than the threshold.

### Optimal splicing from transcript proportions (OSTP)

In each sample we divided the level of each isoform by the sum of levels of the other isoforms to obtain isoform proportions. For each proportion of false positives (FP) we found the t-statistic that maximizes the proportion of true positives (TP) in each simulated dataset, where the proportion of TPs is the proportion of differentially spliced isoforms with a fold change of 2 or greater that are detected. We here refer to the results obtained in this way as optimal splicing from isoform proportions, or OSTP. Note that OSTP is not a method that can be used in practice, since it does not provide a way to choose the t-statistic threshold for a given dataset, unless this is baked into a simulation. Note that OSTP is not a method that can be used in practice, since it does not provide a way to choose the t-statistic threshold for a given dataset, unless this is baked into a simulation.

### MCMC

We used the stan package for MCMC modeling via its R interface rstan. ^38^ We set the number of chains which is set to 1, the number of warmup steps to 2000, and the number of total steps to 10000. In order to obtain an extended range of p-values for generating precision-recall curves in the simulation, we set the number of steps to 100000, and the number of warmup iterations to 10000. We used the R package coda to parse the results.

### GTEx dataset

The genome-wide, cross-tissue expression profiling provided by GTEx includes over 17 thousand expression samples from 948 donors in 54 tissues. ^15^

RSEM ^35^ isoform counts were extracted from the file `GTEx_Analysis_2016-01-15_v7_RSEMv1.2.22_transcript_expected_count.txt`, which is available from the GTEx portal. ^15^ This file contains RSEM counts for multiple isoforms of different genes identified by Ensembl ids, in different tissues of different donors. We used biomaRt to map Ensembl ids to HGNC gene symbols. ^39^ The sample annotations were extracted from the file `GTEx_v7_Annotations_SampleAttributesDS.txt`, which contains data from 8444 samples from 703 donors.

### Robustness analysis of HBA-DEALS

We reasoned that if HBA-DEALS is able to robustly identify mRNAs that are consistently differentially expressed, alternatively spliced, both, or neither, then we should observe a high level of consistency in its results for subsets of samples in the GTEx dataset. Therefore, for each pair of tissues we randomly divided the data into subcohorts of 30 samples, 15 from each condition, keeping transcripts that had a count of at least 1 in each subcohort sample. We then ran HBA-DEAL on each subcohort separately, and compared the sets of genes and isoforms that were identified as differentially expressed and differentially spliced, respectively. There was a highly significant overlap amongst both genes and isoforms that were consistently identified between cohorts. We then compared the changes in gene expression levels and isoform proportions quantitatively, by computing the R-square value between log-fold expression changes of each gene and log-fold isoform proportion changes of each isoform. This has resulted in high correlation between cohorts in the different tissues (figure 2 c-d). As expected, tissues that were not related showed an overall higher correlation, since differences between tissues are much greater than differences between donors.

Fold changes in expression were calculated as the mean of the posterior beta, and fold changes in splicing as the Aitchison perturbation between the mean of the posterior of the fraction in controls and the mean of the posterior of alpha divided by the mean of the posterior of the fraction in controls.

To further determine the robustness of the set of isoforms that were identified as differentially spliced, we converted the GTEx expression data into isoform proportions, and selected the set of isoforms that were differentially spliced in at least 3 cohorts and had a fold change of at least 2 for a multidimensional scaling of all the samples together. We used the R function cmdscale. The clear separation between samples belonging to different tissues confirmed the robustness of the isoforms identified in individual cohorts and their consistency as a set. In order to validate the visual observation, we computed the ratio of the mean between-tissue-distances to within-tissue-distances in the MDS for the real data and for 1000 permutations of the tissue labels, for each pair of tissues. We then counted the number of times that a value computed for the permuted labels was greater or equal to the corresponding value computed for the original labels. For all 10 tissue pairs, this did not occur in any of the permutations, corresponding to a p-value ¡ 0.001 that a separation between two labels occurred by chance.

### Multidimensional scaling

Multidimensional scaling (MDS) is a nonlinear transformation that translates a matrix of pairwise distances between objects into a two-dimensional visualization of the objects that preserves the pairwise distances as much as possible. ^19;40^

### Gene Ontology analysis

For Gene Ontology enrichment analysis we used the program Ontologizer, ^41^ using the Parent-child Intersection algorithm. ^42^ The population set was composed of all the genes that passed the minimum-counts threshold, i.e. that participated in the analysis. The complete lists of GO categories with Bonferroni-corrected *p*-values less or equal to 0.01 are given in supplementary tables S5, S6, S7, and S8. We used the `go.obo` and `goa_human.gaf` files downloaded on December 2, 2019.

### Defining four mRNA categories

We define splicing-regulated genes as genes for which differential splicing was observed at least K times as often as differential expression, and expression-regulated genes as genes for which differential expression was observed at least K times as often as differential splicing of one of the isoforms, K=2. To test for robustness of this criterion, we repeated the analysis for increasing values of K. As can been seen in Supplemental Fig. S3, the properties that were found for K=2 remain significant for increasing values of K.

### Additional data sources

We have used several additional data sources in order to characterize the properties if splicing- and expression-regulated genes. TF targets, the TATA box motif in promoters, and dispersion of the transcription start site (TSS) were obtained from the FANTOM project portal ^43^. For TF targets we used the file `hg38.gencode_v28.TF_HUMAN.tsv`, where TF is the transcription factor name. Gene lengths were retrieved using the biomaRt R package. Exon and isoform annotations were taken from the file `Homo_sapiens.GRCh38.91.gtf` that was download from the Ensembl website. The methylation datasets were downloaded from MethBank ^44^. The age groups were: age0, age2-4, age5-13, age14-16, age17-28, age29-36, age37-42, age43-53, age54-66, age67-75, age76-88, and age89-101.

### Simulation

We used the code provided with ^6^ to simulate isoform levels and followed the methodology that was used to add differential expression for adding differential splicing. For each gene, a number of expressed isoforms was randomly generated between 2 and 10 using the probabilities 0.4,0.2,0.1,0.05,0.05,0.05,0.05,0.05,0.05. The proportions of genes were then divided to by the corresponding number of isoforms. A set of differentially spliced genes was selected at random. In the original code, a gene’s proportion is multiplied by 2 in either cases or controls to generate differential expression. Therefore, for each differentially spliced gene, the proportion of one random isoform was increased 2-fold in cases, and the proportion of another isoform was increased 2-fold in control. This ensures that the total proportion of the gene remains unchanged. After generating isoform proportions, the simulation proceeds the same as the original code. We generated datasets with random seeds 1-50. The first 25 were generated using equal library sizes and the last 25 with unequal library sizes. The input to rMATS consists of counts for “skip” and “inclusion” counts, each representing a distinct isoform. We set for each isoform the “skip” count as the number of counts of the isoform, and the “inclusion” counts the number of counts of the other isoforms. We set isoform length to 1 and the PSI cutoff to 1e-10. Tools provided as R packages were used according to the usage instructions in the packages. In order to generate mean precision-recall curves, we fitted a trendline with the function lowess in the R package limma to the precision and recall values for each tool in each dataset. For missing precision values, we added points with the nearest lower recall value. The mean of the trendline values over all the datasets was then computed for each tool to obtain the mean precision-recall curve.

### Optimal splicing from transcript proportions (OSTP)

The OSTP method obtains the best precision-recall tradeoff possible using a t-test. First, it calculates the t-statistic for the difference between isoform proportions in cases and controls. Then, for each threshold value of the statistic, it finds the false positive (FP) proportion and true positive (TP) proportion. Finally, for each FP proportion it selects the optimal TP proportion.

### Statistical tests

For performing the Mann-Whitney test, Fisher’s Exact Test, creating the logistic regression model, computing the hypergeometric cdf, the t statistic and the Mann-Whitney statistic we used the core modules R programming language.

### Dispersion

Promoters can be characterized as either sharp type or broad type, depending on whether they contain one dominant transcription start site or multiple transcription start sites. ^45^ Cap analysis of gene expression (CAGE) can be used to identify transcription start sites in promoters. CAGE experiments generate sets of 20 to 27 bp sequence tags from the 5’ ends of mRNA, which can be matched to a reference genome. Any accumulation of tags (“peak”) is a reliable indicator of a transcription start site.

Based on FANTOM5 data, ^46^ we computed dispersion indexes of CAGE tags for all promoter sequences, a metric that is conceptually similar to the standard deviation of tag counts. ^47^ A low dispersion index indicates a sharp distribution of tags, and a high dispersion index indicates a broad distribution of tags. To compute dispersion indexes, we counted tags between positions −99 and +100 relative to and on the same strand as the annotated transcription start sites. Let *s* be the dispersion index and *x_i_* be the number of tags at position *i*. Then,

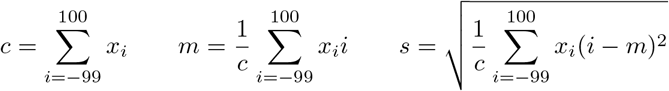

The significance of the difference between the the DAST and the DGE groups was determined using the Mann-Whitney test.

### CpG islands

In the human genome, CpG dinucleotides are present at about 20% of the frequency that would be expected based on the overall GC-content. The depletion of CpG dinucleotides in the human and other mammalian genomes is due to the increased mutability of methylcytosine within CpG dinucleotides. Stretches of GC-rich (~65%) sequence in which the observed frequency of CpG dinucleotides is close to the frequency that would be expected based on the individual frequency of G and C bases are termed CpG islands (CGIs). CGIs are associated with the upstream region of many genes generally covering all or part of the promoter and typically display an average size of about 1 kb ^48;49^.

To identify CGIs in this study, a 100-nucleotide window was shifted in 1 bp intervals across the promoter sequences from position [−200, −100) relative to the TSS to [+100, +200). The percentage GC-content and CpG expected/observed ratio

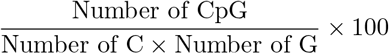

were calculated per window.

A promoter was considered having a CGI if the consecutive windows inside any region spanning at least 200 nt all had GC-contents ≥ 50% and CpG expected/observed ratios ≥ 0.6^50^.

The significance of the difference between the DAST and DGE groups was determined using the Mann-Whitney test.

### TATA box

We employed a matrix of counts for TATA to define a position count matrix (PCM). ^51^

A corresponding position weight matrix (PWM) was computed. A PWM of length *ℓ* assigns each oligonucleotide of length *ℓ* a matching score 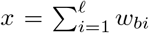, where *w_bi_* is the weight of base *b* at column *i* of the matrix. The weights *w_bi_* were computed relative to the log-normalized base frequencies per position of the PCM. We identified TATA boxes in the window [−32, −28] with respect to the transcription start site if the PWM matching score of any oligonucleotide beginning in this region exceeded a threshold of 0.790. ^51^

The significance of the difference between the DAST and DGE groups was determined using the Mann-Whitney test.

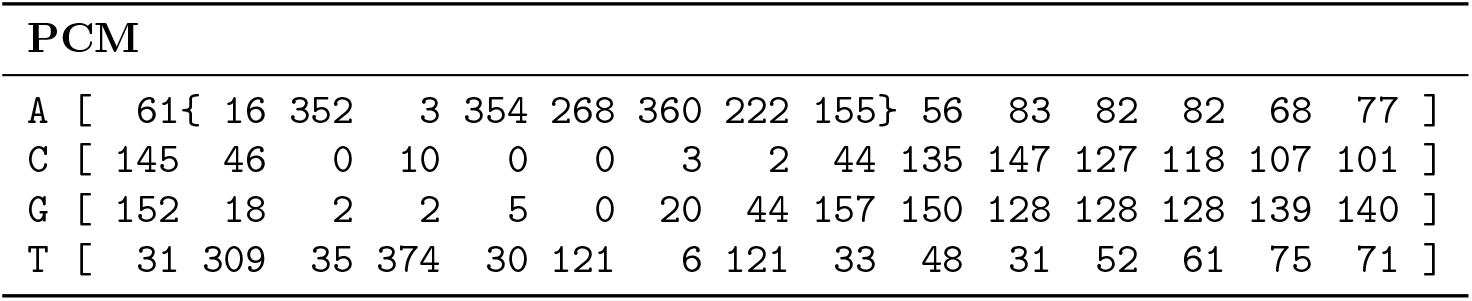

### Number of isoforms

The number of isoforms per gene was retrieved from the GTF file “Homo sapiens.GRCh38.91.gtf”. The significance of the difference between the DAST and DGE groups was determined using the Mann-Whitney test.

### Exon length

Exon lengths were retrieved from the GTF file “Homo sapiens.GRCh38.91.gtf”. The significance of the difference between the DAST and DGE groups was determined using the Mann-Whitney test.

### Software

HBA-DEALS is implemented as an R package that is freely available at https://github.com/TheJacksonLaboratory/HBA-DEALS.

## References

[1] Pollard, M. O., Gurdasani, D., Mentzer, A. J., Porter, T., & Sandhu, M. S. Long reads: their purpose and place. Human molecular genetics 27:R234–R241 (2018).

[2] Wang, Z., Gerstein, M., & Snyder, M. RNA-seq: a revolutionary tool for transcriptomics. Nature reviews. Genetics 10:57–63 (2009).

[3] Stark, R., Grzelak, M., & Hadfield, J. RNA sequencing: the teenage years. Nature reviews. Genetics 20:631–656 (2019).

[4] Robinson, M. D., McCarthy, D. J., & Smyth, G. K. edgeR: a Bioconductor package for differential expression analysis of digital gene expression data. Bioinformatics (Oxford, England) 26:139–140 (2010).

[5] Love, M. I., Huber, W., & Anders, S. Moderated estimation of fold change and dispersion for RNA-seq data with DESeq2. Genome biology 15:550 (2014).

[6] Law, C. W., Chen, Y., Shi, W., & Smyth, G. K. voom: Precision weights unlock linear model analysis tools for rna-seq read counts. Genome biology 15:R29 (2014).

[7] Sterne-Weiler, T., Weatheritt, R. J., Best, A. J., Ha, K. C., & Blencowe, B. J. Efficient and accurate quantitative profiling of alternative splicing patterns of any complexity on a laptop. Molecular Cell 72(1):187–200.e6 (2018).

[8] Shen, S., et al. rMATS: robust and flexible detection of differential alternative splicing from replicate RNA-Seq data. Proceedings of the National Academy of Sciences of the United States of America 111:E5593–E5601 (2014).

[9] Katz, Y., Wang, E. T., Airoldi, E. M., & Burge, C. B. Analysis and design of RNA sequencing experiments for identifying isoform regulation. Nature Methods 7(12):1009–1015 (2010).

[10] Hu, Y., et al. DiffSplice: the genome-wide detection of differential splicing events with RNA-seq. Nucleic Acids Research 41(2):e39–e39 (2012).

[11] Sebestyén, E., Zawisza, M., & Eyras, E. Detection of recurrent alternative splicing switches in tumor samples reveals novel signatures of cancer. Nucleic Acids Research 43(3):1345–1356 (2015).

[12] Kahles, A., Ong, C. S., Zhong, Y., & Rätsch, G. SplAdder: identification, quantification and testing of alternative splicing events from RNA-seq data. Bioinformatics 32(12):1840–1847 (2016).

[13] Climente-González, H., Porta-Pardo, E., Godzik, A., & Eyras, E. The functional impact of alternative splicing in cancer. Cell Reports 20(9):2215–2226 (2017).

[14] Oshlack, A. & Wakefield, M. J. Transcript length bias in rna-seq data confounds systems biology. Biology direct 4:14 (2009).

[15] GTEx Consortium. The genotype-tissue expression (GTEx) project. Nature genetics 45:580–585 (2013).

[16] Aitchison, J. The Statistical Analysis of Compositional Data. Springer Netherlands (1986).

[17] Hardcastle, T. J. & Kelly, K. A. baySeq: empirical Bayesian methods for identifying differential expression in sequence count data. BMC bioinformatics 11:422 (2010).

[18] Tarazona, S., et al. Data quality aware analysis of differential expression in RNA-seq with NOISeq R/Bioc package. Nucleic acids research 43:e140 (2015).

[19] Mardia, K. Some properties of clasical multi-dimesional scaling. Communications in Statistics - Theory and Methods 7(13):1233–1241 (1978).

[20] The Gene Ontology Consortium. Expansion of the Gene Ontology knowledgebase and resources. Nucleic acids research 45:D331–D338 (2017).

[21] Fu, X.-D. & Ares, M. Context-dependent control of alternative splicing by RNA-binding proteins. Nature reviews. Genetics 15:689–701 (2014).

[22] Pimentel, H., et al. A dynamic intron retention program enriched in rna processing genes regulates gene expression during terminal erythropoiesis. Nucleic acids research 44:838–851 (2016).

[23] Rodríguez, S. A., et al. Global genome splicing analysis reveals an increased number of alternatively spliced genes with aging. Aging cell 15:267–278 (2016).

[24] Shirai, C. L., et al. Mutant u2af1 expression alters hematopoiesis and pre-mrna splicing in vivo. Cancer cell 27:631–643 (2015).

[25] Young, J. I., et al. Regulation of rna splicing by the methylation-dependent transcriptional repressor methyl-cpg binding protein 2. Proceedings of the National Academy of Sciences of the United States of America 102:17551–17558 (2005).

[26] Shukla, S., et al. Ctcf-promoted rna polymerase ii pausing links dna methylation to splicing. Nature 479:74–79 (2011).

[27] Lev Maor, G., Yearim, A., & Ast, G. The alternative role of DNA methylation in splicing regulation. Trends in genetics: TIG 31:274–280 (2015).

[28] Cramer, P., et al. Coupling of transcription with alternative splicing: RNA pol ii promoters modulate SF2/ASF and 9G8 effects on an exonic splicing enhancer. Molecular cell 4:251–258 (1999).

[29] Damgaard, C. K., et al. A 5’ splice site enhances the recruitment of basal transcription initiation factors in vivo. Molecular cell 29:271–278 (2008).

[30] Giurgiu, M., et al. Corum: the comprehensive resource of mammalian protein complexes-2019. Nucleic acids research 47:D559–D563 (2019).

[31] Malygin, A. A., Parakhnevitch, N. M., Ivanov, A. V., Eperon, I. C., & Karpova, G. G. Human ribosomal protein s13 regulates expression of its own gene at the splicing step by a feedback mechanism. Nucleic acids research 35:6414–6423 (2007).

[32] Takei, S., Togo-Ohno, M., Suzuki, Y., & Kuroyanagi, H. Evolutionarily conserved autoregulation of alternative pre-mrna splicing by ribosomal protein l10a. Nucleic acids research (2016).

[33] Lareau, L. F. & Brenner, S. E. Regulation of splicing factors by alternative splicing and nmd is conserved between kingdoms yet evolutionarily flexible. Molecular biology and evolution 32:1072–1079 (2015).

[34] Ravasi, T., et al. An atlas of combinatorial transcriptional regulation in mouse and man. Cell 140:744–752 (2010).

[35] Li, B. & Dewey, C. N. RSEM: accurate transcript quantification from RNA-seq data with or without a reference genome. BMC bioinformatics 12:323 (2011).

[36] Pertea, M., et al. StringTie enables improved reconstruction of a transcriptome from RNA-seq reads. Nature Biotechnology 33:290–295 (2015).

[37] Tardaguila, M., et al. SQANTI: extensive characterization of long-read transcript sequences for quality control in full-length transcriptome identification and quantification. Genome research [Epub ahead of print] (2018).

[38] Carpenter, B., et al. Stan: A probabilistic programming language. Journal of Statistical Software 76(1) (2017).

[39] Smedley, D., et al. Biomart–biological queries made easy. BMC genomics 10:22 (2009).

[40] Hout, M. C., Papesh, M. H., & Goldinger, S. D. Multidimensional scaling. Wiley interdisciplinary reviews. Cognitive science 4:93–103 (2013).

[41] Bauer, S., Grossmann, S., Vingron, M., & Robinson, P. N. Ontologizer 2.0–a multifunctional tool for GO term enrichment analysis and data exploration. Bioinformatics (Oxford, England) 24:1650–1651 (2008).

[42] Grossmann, S., Bauer, S., Robinson, P. N., & Vingron, M. Improved detection of overrepresentation of Gene-Ontology annotations with parent child analysis. Bioinformatics (Oxford, England) 23:3024–3031 (2007).

[43] Noguchi, S., et al. FANTOM5 CAGE profiles of human and mouse samples. Scientific data 4:170112 (2017).

[44] Li, R., et al. MethBank 3.0: a database of DNA methylomes across a variety of species. Nucleic acids research 46:D288–D295 (2018).

[45] Carninci, P., et al. Genome-wide analysis of mammalian promoter architecture and evolution. Nat Genet 38(6):626–635 (2006).

[46] Arner, E., et al. Transcribed enhancers lead waves of coordinated transcription in transitioning mammalian cells. Science 347(6225):1010–1014 (2015).

[47] Dreos, R., Ambrosini, G., & Bucher, P. Influence of rotational nucleosome positioning on transcription start site selection in animal promoters. PLoS computational biology 12:e1005144 (2016).

[48] Larsen, F., Gundersen, G., Lopez, R., & Prydz, H. Cpg islands as gene markers in the human genome. Genomics 13:1095–1107 (1992).

[49] Robinson, P. N., Böhme, U., Lopez, R., Mundlos, S., & Nürnberg, P. Gene-ontology analysis reveals association of tissue-specific 5’ cpg-island genes with development and embryogenesis. Human molecular genetics 13:1969–1978 (2004).

[50] Gardiner-Garden, M. & Frommer, M. Cpg islands in vertebrate genomes. Journal of molecular biology 196:261–282 (1987).

[51] Bucher, P. Weight matrix descriptions of four eukaryotic rna polymerase ii promoter elements derived from 502 unrelated promoter sequences. Journal of molecular biology 212:563–578 (1990).

