## Supplemental material for "Accurate and simultaneous identification of differential expression and splicing using hierarchical Bayesian analysis"

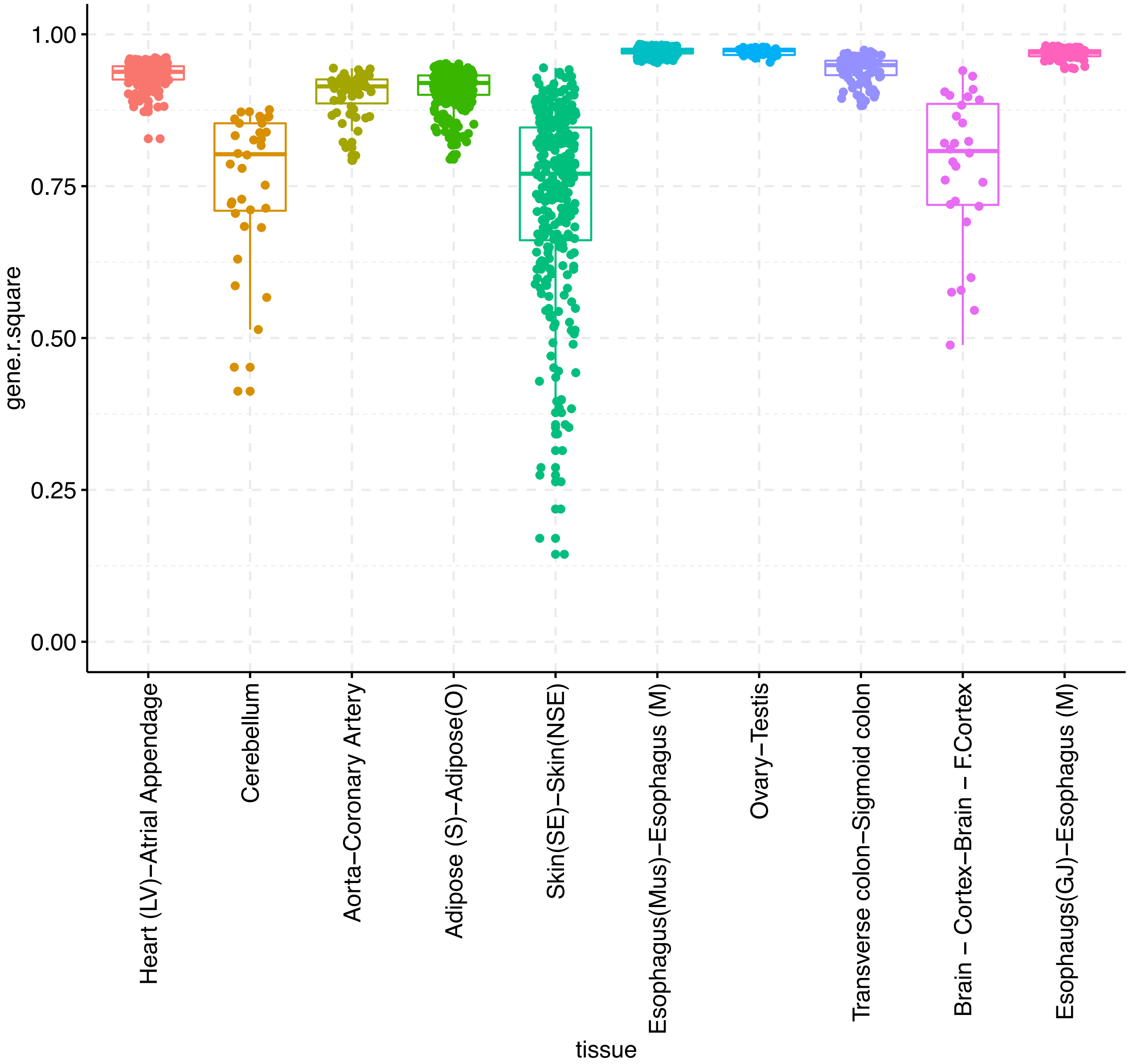


**Figure S1:** Square of the correlations ($R^2$) of gene expression in comparisons between 'Near" tissues as defined in Table S1.

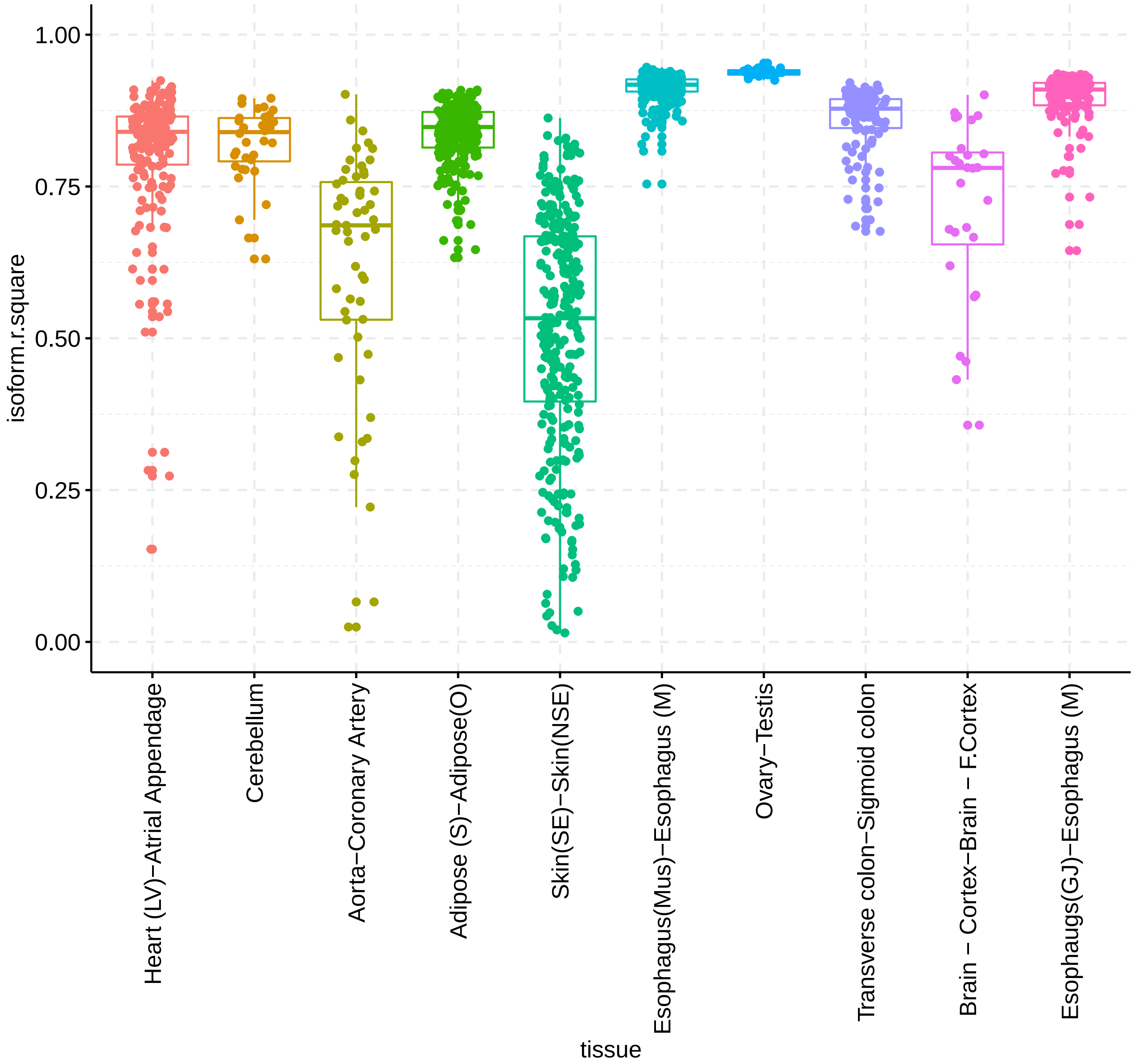


**Figure S2:** Square of the correlations ($R^2$) of isoform proportions in comparisons between 'Near" tissues as defined in Table S1.

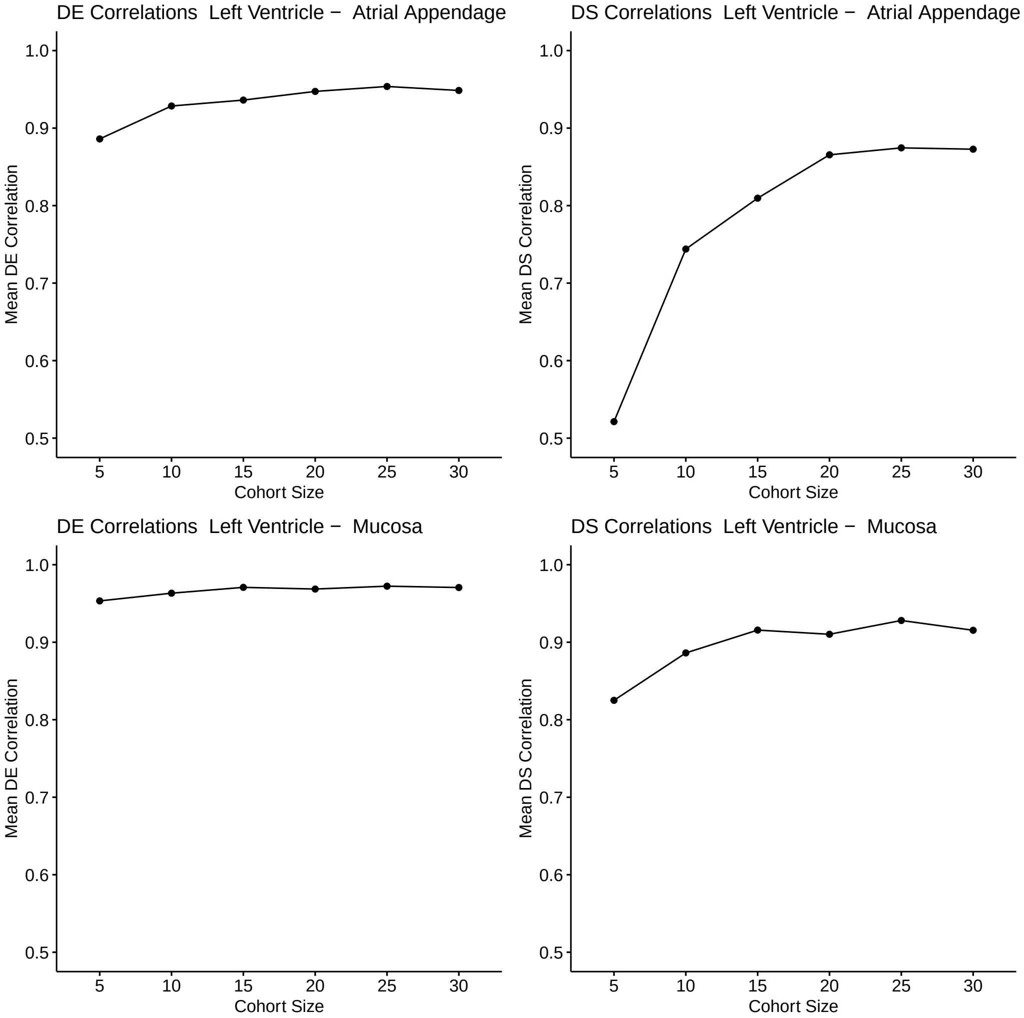


**Figure S3:** Influence of sample size on $R^2$ values for correlations of differential gene expression (DE) and differential splicing (DS). Analysis was performed as with the analysis shown in Figure. 2c–d of the main manuscript and Figures S1 and S2.

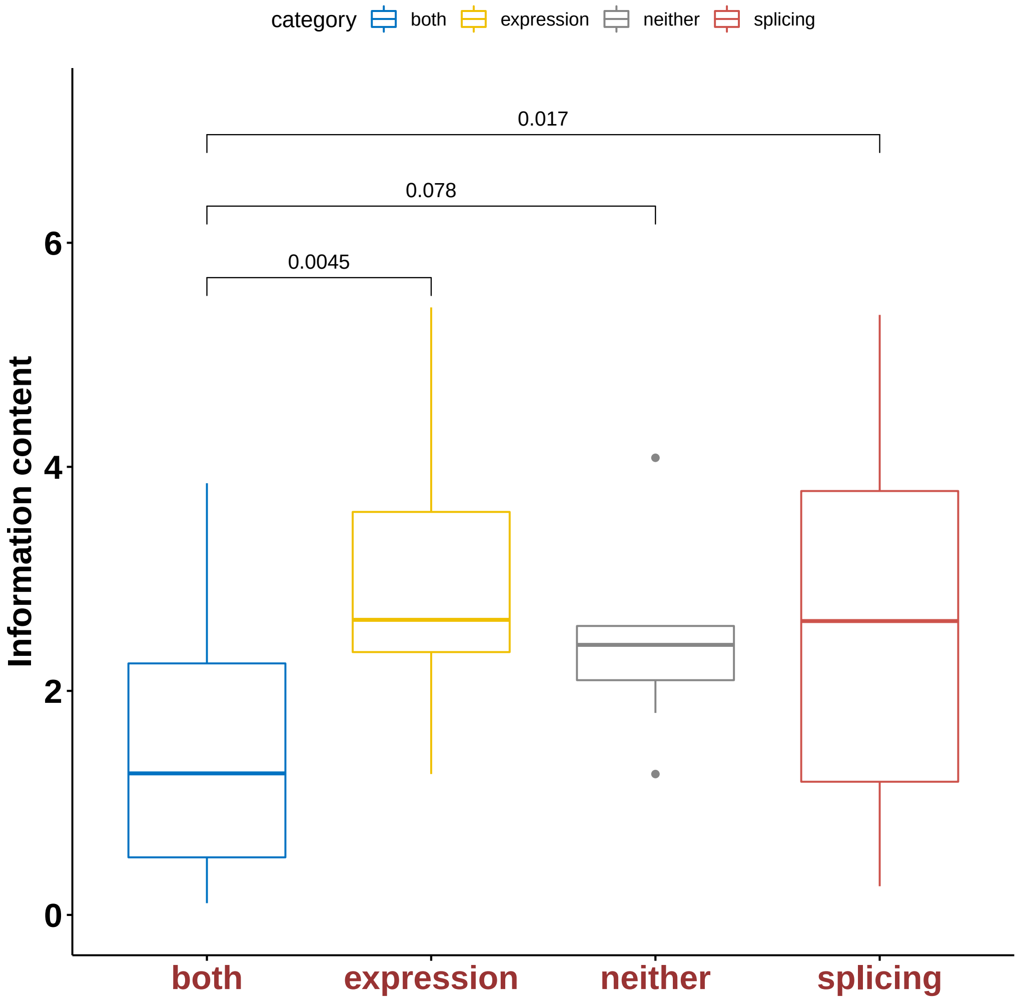


**Figure S4:** The information content of the GO terms that annotate the gene groups with preferential alternative splicing (splicing, Table S5, preferential differential expression (expression, Table S6, both differential alternative splicing and differential expression (both, Table S7, and neither differential alternative splicing nor differential gene expression (neither, Table S8) are shown. Means were compared with a two-sided t-test. "Both" had the lowest mean information content. The two groups "splicing" and "expression" had significantly higher mean information content values.

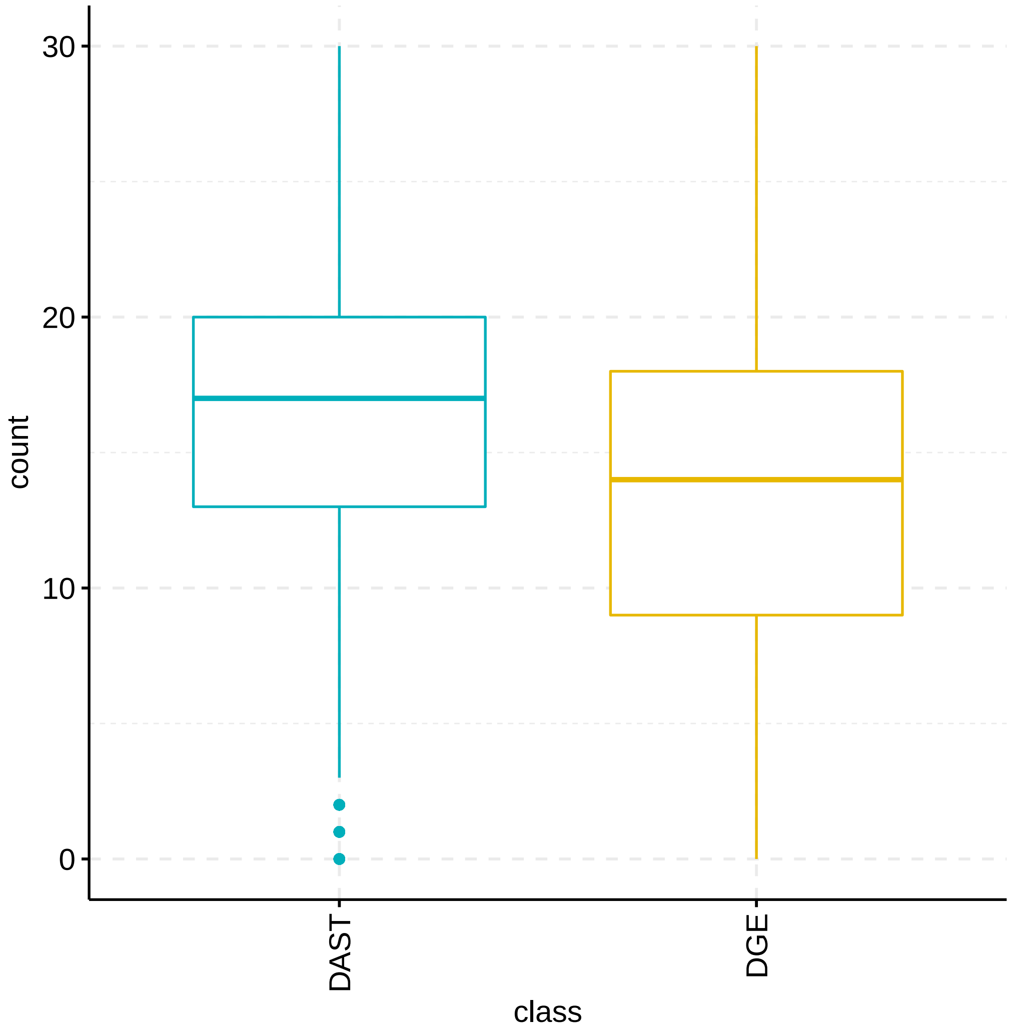


**Figure S5:** Counts of DAST transcription factors (TFs) with TF binding motifs (TFBMs) in promoters of all DAST genes and the corresponding count with TFBMs in promoters of DGE genes. There was a mea of 16.05 in the DAST genes vs 13.67 in the DGE genes, $p = 7.36 \times 10^{-53}$, Mann-Whitney test.

| Tissue1 | Tissue2 | Type | Comparisons (n) |
|---|---|---|---|
| Heart - Left Ventricle Heart | Atrial Appendage | N | 171 |
| Brain - Cerebellum | Brain - Cerebellar Hemisphere | N | 36 |
| Artery - Aorta | Artery - Coronary | N | 55 |
| Adipose - Subcutaneous | Adipose - Visceral (Omentum) | N | 253 |
| Skin - Sun Exposed (Lower leg) | Skin - Not Sun Exposed (Suprapubic) | N | 300 |
| Esophagus - Muscularis | Esophagus - Mucosa | N | 276 |
| Ovary | Testis | N | 28 |
| Colon - Transverse | Colon - Sigmoid | N | 105 |
| Brain - Cortex | Brain - Frontal Cortex (BA9) | N | 28 |
| Esophagus - Gastroesophageal Junction | Esophagus - Mucosa | N | 120 |
| Heart - Left Ventricle | Esophagus - Mucosa | D | 190 |
| Artery - Coronary | Colon - Transverse | D | 55 |
| Ovary | Brain - Frontal Cortex (BA9) | D | 28 |
| Skin - Sun Exposed (Lower leg) | Colon - Sigmoid | D | 105 |
| Esophagus - Muscularis | Brain - Cerebellum | D | 55 |
| Muscle - Skeletal | Lung | D | 378 |
| Nerve - Tibial | Thyroid | D | 351 |
| Breast - Mammary Tissue | Stomach | D | 136 |
| Pancreas | Adrenal Gland | D | 66 |
| Liver | Pituitary | D | 55 |

**Table S1:** Comparisons between cohorts in GTEx. For each tissue type, 15 samples were chosen at random. Pairs of cohorts were then compared for the analysis described in the main manuscript. The column *Type* indicates whether the compared tissues are considered Near (N) or Distant (D).

| Tissue | Abbreviation |
|---|---|
| Heart - Left Ventricle-Esophagus - Mucosa | H-LV-M |
| Artery - Coronary-Colon - Transverse | A-C-T |
| Ovary-Brain - Frontal Cortex (BA9) | O-FCB |
| Skin - Sun Exposed (Lower leg)-Colon - Sigmoid | S-SELl-S |
| Esophagus - Muscularis-Brain - Cerebellum | E-M-C |
| Muscle - Skeletal-Lung | M-S |
| Nerve - Tibial-Thyroid | N-T |
| Breast - Mammary Tissue-Stomach | B-MT |
| Pancreas-Adrenal Gland | P-AG |
| Liver-Pituitary | Lv-P |

**Table S2:** Abbreviations used in the main manuscript for GTEx tissue types.

| Term | DAST | DGE | DAST/DGE | static | population |
|---|---|---|---|---|---|
| ncRNA metabolic process (GO:0034660) | 113/1757 (6.43%) | 64/3146 (2.03%) | 232/6967 (3.33%) | 15/1804 (0.83%) | 424/13667 (3.10%) |
| RNA processing (GO:0006396) | 234/1757 (13.32%) | 88/3146 (2.80%) | 410/6967 (5.88%) | 25/1804 (1.39%) | 756/13667 (5.53%) |
| RNA binding (GO:0003723) | 373/1757 (21.23%) | 146/3146 (4.64%) | 790/6967 (11.34%) | 70/1804 (3.88%) | 1377/13667 (10.08%) |
| ribonucleoprotein complex subunit organization (GO:0071826) | 169/1757 (9.62%) | 56/3146 (1.78%) | 392/6967 (5.63%) | 35/1804 (1.94%) | 650/13667 (4.76%) |
| mRNA metabolic process (GO:0016071) | 215/1757 (12.24%) | 62/3146 (1.97%) | 375/6967 (5.38%) | 26/1804 (1.44%) | 676/13667 (4.95%) |
| ribonucleoprotein complex (GO:1990904) | 221/1757 (12.58%) | 68/3146 (2.16%) | 405/6967 (5.81%) | 35/1804 (1.94%) | 727/13667 (5.32%) |

**Table S3:** Details for GO terms shown in Fig. 3b of the main manuscript. These terms were chosen to have significant overrepresentation in each one study set with at least 20 annotated genes, whereby there was an at least 2-fold enrichment compared to the popultion set. Details are found in Tables S5–S8.

**(a)** Differential alternative splicing (n=1757)

| **Term** | | **Population** | **Study** | ***p*-value** |
|---|---|---|---|---|
| GO:0032991 | protein-containing complex | 4227 (30.9%) | 791 (45.0%) | $4.35 \times 10^{-31}$ |
| GO:0031974 | membrane-enclosed lumen | 4612 (33.7%) | 820 (46.7%) | $4.48 \times 10^{-24}$ |
| GO:0016071 | mRNA metabolic process | 732 (5.4%) | 238 (13.5%) | $3.04 \times 10^{-22}$ |
| GO:0006396 | RNA processing | 776 (5.7%) | 244 (13.9%) | $3.25 \times 10^{-20}$ |
| GO:0003723 | RNA binding | 1387 (10.1%) | 377 (21.5%) | $7.77 \times 10^{-19}$ |
| GO:0051641 | cellular localization | 2438 (17.8%) | 412 (23.4%) | $1.06 \times 10^{-13}$ |
| GO:0005622 | intracellular | 10834 (79.3%) | 1544 (87.9%) | $1.23 \times 10^{-13}$ |
| GO:0043170 | macromolecule metabolic process | 6708 (49.1%) | 1091 (62.1%) | $6.85 \times 10^{-13}$ |
| GO:1990904 | ribonucleoprotein complex | 585 (4.3%) | 186 (10.6%) | $8.91 \times 10^{-13}$ |
| GO:0070647 | protein modification by small protein conjugation or removal | 858 (6.3%) | 208 (11.8%) | $1.11 \times 10^{-11}$ |
| GO:0022613 | ribonucleoprotein complex biogenesis | 363 (2.7%) | 118 (6.7%) | $1.43 \times 10^{-11}$ |
| GO:0051276 | chromosome organization | 881 (6.4%) | 209 (11.9%) | $6.51 \times 10^{-11}$ |
| GO:0005634 | nucleus | 5577 (40.8%) | 977 (55.6%) | $7.59 \times 10^{-11}$ |

**(b)** Differential gene expression (n=3147)

| **Term** | | **Population** | **Study** | ***p*-value** |
|---|---|---|---|---|
| GO:0031224 | intrinsic component of membrane | 3208 (23.5%) | 881 (28.0%) | $2.66 \times 10^{-14}$ |

**(c)** Differential alternative splicing and gene expression (n=6967)

| **Term** | | **Population** | **Study** | ***p*-value** |
|---|---|---|---|---|
| GO:0005622 | intracellular | 10834 (79.3%) | 5921 (85.0%) | $3.97 \times 10^{-29}$ |
| GO:0043226 | organelle | 9986 (73.1%) | 5508 (79.1%) | $1.21 \times 10^{-25}$ |

**(d)** Static (n=1804)

| **Term** | | **Population** | **Study** | ***p*-value** |
|---|---|---|---|---|
| GO:0031224 | intrinsic component of membrane | 3208 (23.5%) | 512 (28.4%) | $1.79 \times 10^{-14}$ |

**Table S4:** Gene Ontology (GO) terms characterizing **(a)** genes that were preferentially alternatively spliced in the GTEx comparisons, **(b)** genes that were preferentially differentially expressed, **(c)** genes that were both alternatively spliced and differentially expressed, and **(d)** genes that were neither alternatively spliced nor differentially expressed. Significant GO terms with a $p$-value less than $10^{-10}$ are shown. Full results are given in Supplemental Tables TODO. Overrepresentation analysis by the Parent-Child Intersection method of the Ontologizer.[1,2] The population set, defined as the union of the four sets, contained 13,668 genes.

| Term | | Population | Study | p-value |
|---|---|---|---|---|
| GO:0032991 | protein-containing complex | 4227 (30.9%) | 791 (45.0%) | $4.35 \times 10^{-31}$ |
| GO:0031974 | membrane-enclosed lumen | 4612 (33.7%) | 820 (46.7%) | $4.48 \times 10^{-24}$ |
| GO:0016071 | mRNA metabolic process | 732 (5.4%) | 238 (13.5%) | $3.04 \times 10^{-22}$ |
| GO:0006396 | RNA processing | 776 (5.7%) | 244 (13.9%) | $3.25 \times 10^{-20}$ |
| GO:0003723 | RNA binding | 1387 (10.1%) | 377 (21.5%) | $7.77 \times 10^{-19}$ |
| GO:0051641 | cellular localization | 2438 (17.8%) | 412 (23.4%) | $1.06 \times 10^{-13}$ |
| GO:0005622 | intracellular | 10834 (79.3%) | 1544 (87.9%) | $1.23 \times 10^{-13}$ |
| GO:0043170 | macromolecule metabolic process | 6708 (49.1%) | 1091 (62.1%) | $6.85 \times 10^{-13}$ |
| GO:1990904 | ribonucleoprotein complex | 585 (4.3%) | 186 (10.6%) | $8.91 \times 10^{-13}$ |
| GO:0070647 | protein modification by small protein conjugation or removal | 858 (6.3%) | 208 (11.8%) | $1.11 \times 10^{-11}$ |
| GO:0022613 | ribonucleoprotein complex biogenesis | 363 (2.7%) | 118 (6.7%) | $1.43 \times 10^{-11}$ |
| GO:0051276 | chromosome organization | 881 (6.4%) | 209 (11.9%) | $6.51 \times 10^{-11}$ |
| GO:0005634 | nucleus | 5577 (40.8%) | 977 (55.6%) | $7.59 \times 10^{-11}$ |
| GO:0003676 | nucleic acid binding | 3428 (25.1%) | 656 (37.3%) | $6.89 \times 10^{-10}$ |
| GO:0043226 | organelle | 9986 (73.1%) | 1438 (81.8%) | $1.46 \times 10^{-9}$ |
| GO:1901363 | heterocyclic compound binding | 4795 (35.1%) | 801 (45.6%) | $1.81 \times 10^{-9}$ |
| GO:0097159 | organic cyclic compound binding | 4837 (35.4%) | 806 (45.9%) | $2.65 \times 10^{-9}$ |
| GO:0033036 | macromolecule localization | 2683 (19.6%) | 424 (24.1%) | $2.68 \times 10^{-8}$ |
| GO:0019787 | ubiquitin-like protein transferase activity | 424 (3.1%) | 106 (6.0%) | $9.42 \times 10^{-8}$ |
| GO:0071705 | nitrogen compound transport | 2102 (15.4%) | 352 (20.0%) | $3.14 \times 10^{-7}$ |
| GO:0008152 | metabolic process | 8160 (59.7%) | 1214 (69.1%) | $3.28 \times 10^{-7}$ |
| GO:0101005 | ubiquitinyl hydrolase activity | 220 (1.6%) | 64 (3.6%) | $2.65 \times 10^{-6}$ |
| GO:0140098 | catalytic activity, acting on RNA | 344 (2.5%) | 86 (4.9%) | $3.84 \times 10^{-6}$ |
| GO:0071826 | ribonucleoprotein complex subunit organization | 169 (1.2%) | 64 (3.6%) | $3.92 \times 10^{-6}$ |
| GO:0006403 | RNA localization | 206 (1.5%) | 66 (3.8%) | $7.88 \times 10^{-6}$ |
| GO:0043933 | protein-containing complex subunit organization | 1645 (12.0%) | 309 (17.6%) | $3.55 \times 10^{-5}$ |
| GO:0033554 | cellular response to stress | 1606 (11.8%) | 263 (15.0%) | $5.93 \times 10^{-5}$ |
| GO:0010608 | posttranscriptional regulation of gene expression | 526 (3.8%) | 135 (7.7%) | $7.17 \times 10^{-5}$ |
| GO:0120114 | Sm-like protein family complex | 135 (1.0%) | 53 (3.0%) | $9.4 \times 10^{-5}$ |
| GO:0005829 | cytosol | 4135 (30.3%) | 663 (37.7%) | 0.000117 |
| GO:0044419 | interspecies interaction between organisms | 751 (5.5%) | 160 (9.1%) | 0.000251 |
| GO:0046483 | heterocycle metabolic process | 4420 (32.3%) | 752 (42.8%) | 0.000304 |
| GO:0005840 | ribosome | 218 (1.6%) | 65 (3.7%) | 0.000555 |
| GO:0035770 | ribonucleoprotein granule | 187 (1.4%) | 50 (2.8%) | 0.000565 |
| GO:0019783 | ubiquitin-like protein-specific protease activity | 305 (2.2%) | 81 (4.6%) | 0.000645 |
| GO:0010467 | gene expression | 4000 (29.3%) | 727 (41.4%) | 0.000931 |
| GO:0006725 | cellular aromatic compound metabolic process | 4445 (32.5%) | 751 (42.7%) | 0.00159 |
| GO:0000974 | Prp19 complex | 86 (0.6%) | 36 (2.0%) | 0.00349 |
| GO:0003682 | chromatin binding | 421 (3.1%) | 95 (5.4%) | 0.00665 |
| GO:0032606 | type I interferon production | 101 (0.7%) | 25 (1.4%) | 0.00949 |

**Table S5:** Overrepresentation analysis by the Parent-Child Intersection method of the Ontologizer.[1,2]. Genes displaying preferential alternative splicing ($n = 1757$ genes). Population as in Table 1 of main manuscript ($n = 13,668$ genes). Bonferroni corrected $p$-value is shown, with threshold of 0.01.

| Term | | Population | Study | p-value |
|---|---|---|---|---|
| GO:0031224 | intrinsic component of membrane | 3208 (23.5%) | 881 (28.0%) | $2.66 \times 10^{-14}$ |
| GO:0003677 | DNA binding | 2235 (16.4%) | 491 (15.6%) | $1.41 \times 10^{-10}$ |
| GO:0060089 | molecular transducer activity | 652 (4.8%) | 218 (6.9%) | $6.19 \times 10^{-9}$ |
| GO:0018212 | peptidyl-tyrosine modification | 266 (1.9%) | 82 (2.6%) | $4.42 \times 10^{-5}$ |
| GO:0010876 | lipid localization | 272 (2.0%) | 86 (2.7%) | $5.53 \times 10^{-5}$ |
| GO:0005581 | collagen trimer | 65 (0.5%) | 31 (1.0%) | 0.000333 |
| GO:0009986 | cell surface | 525 (3.8%) | 169 (5.4%) | 0.000388 |
| GO:0051674 | localization of cell | 1122 (8.2%) | 310 (9.9%) | 0.000493 |
| GO:0040011 | locomotion | 1285 (9.4%) | 357 (11.3%) | 0.000596 |
| GO:0071692 | protein localization to extracellular region | 392 (2.9%) | 104 (3.3%) | 0.000634 |
| GO:0001067 | regulatory region nucleic acid binding | 1446 (10.6%) | 324 (10.3%) | 0.00137 |
| GO:0030545 | receptor regulator activity | 206 (1.5%) | 81 (2.6%) | 0.00158 |
| GO:0022610 | biological adhesion | 988 (7.2%) | 278 (8.8%) | 0.00662 |

**Table S6:** Overrepresentation analysis by the Parent-Child Intersection method of the Ontologizer.[1,2]. Genes displaying preferential differential expression ($n = 3147$ genes). Population as in Table 1 of main manuscript ($n = 13,668$ genes). Bonferroni corrected $p$-value is shown, with threshold of 0.01.

| Term | | Population | Study | p-value |
|---|---|---|---|---|
| GO:0005622 | intracellular | 10834 (79.3%) | 5921 (85.0%) | $3.97 \times 10^{-29}$ |
| GO:0043226 | organelle | 9986 (73.1%) | 5508 (79.1%) | $1.21 \times 10^{-25}$ |
| GO:0005737 | cytoplasm | 8761 (64.1%) | 4935 (70.8%) | $2.33 \times 10^{-8}$ |
| GO:0003824 | catalytic activity | 5651 (41.3%) | 3176 (45.6%) | $1.63 \times 10^{-5}$ |
| GO:0030055 | cell-substrate junction | 376 (2.8%) | 259 (3.7%) | $3.18 \times 10^{-5}$ |
| GO:0044281 | small molecule metabolic process | 1544 (11.3%) | 944 (13.5%) | 0.000231 |
| GO:0031974 | membrane-enclosed lumen | 4612 (33.7%) | 2583 (37.1%) | 0.000257 |
| GO:0008152 | metabolic process | 8160 (59.7%) | 4477 (64.3%) | 0.000565 |
| GO:0070161 | anchoring junction | 506 (3.7%) | 330 (4.7%) | 0.00125 |
| GO:0003674 | molecular_function | 11739 (85.9%) | 6263 (89.9%) | 0.0014 |
| GO:0043230 | extracellular organelle | 1579 (11.6%) | 967 (13.9%) | 0.00478 |
| GO:0005829 | cytosol | 4135 (30.3%) | 2444 (35.1%) | 0.00483 |
| GO:0019899 | enzyme binding | 1961 (14.3%) | 1168 (16.8%) | 0.00747 |
| GO:0055114 | oxidation-reduction process | 897 (6.6%) | 560 (8.0%) | 0.00865 |

**Table S7:** Overrepresentation analysis by the Parent-Child Intersection method of the Ontologizer.[1,2]. Genes displaying both alternative splicing and differential expression ($n = 6967$ genes). Population as in Table 1 of main manuscript ($n = 13,668$ genes). Bonferroni corrected $p$-value is shown, with threshold of 0.01.

| Term | | Population | Study | p-value |
|---|---|---|---|---|
| GO:0031224 | intrinsic component of membrane | 3208 (23.5%) | 512 (28.4%) | $1.79 \times 10^{-14}$ |
| GO:0006811 | ion transport | 1224 (9.0%) | 194 (10.8%) | $4.34 \times 10^{-9}$ |
| GO:0055085 | transmembrane transport | 1223 (8.9%) | 185 (10.3%) | $2.63 \times 10^{-6}$ |
| GO:1990351 | transporter complex | 210 (1.5%) | 47 (2.6%) | $8.44 \times 10^{-6}$ |
| GO:0003677 | DNA binding | 2235 (16.4%) | 258 (14.3%) | $1.5 \times 10^{-5}$ |
| GO:0050877 | nervous system process | 624 (4.6%) | 118 (6.5%) | 0.00118 |
| GO:0005215 | transporter activity | 996 (7.3%) | 163 (9.0%) | 0.006 |

**Table S8:** Overrepresentation analysis by the Parent-Child Intersection method of the Ontologizer. [1,2]. Genes displaying neither alternative splicing nor differential expression ($n = 1804$ genes). Population as in Table 1 of main manuscript ($n = 13,668$ genes). Bonferroni corrected $p$-value is shown, with threshold of 0.01.

| Age group | Promoter | Gene body |
|---|---|---|
| 0 y | $1.698 \times 10^{-23}$ | $5.146 \times 10^{-26}$ |
| 2-4 y | $5.754 \times 10^{-26}$ | $4.177 \times 10^{-28}$ |
| 5-13 y | $1.367 \times 10^{-26}$ | $3.163 \times 10^{-28}$ |
| 14-16 y | $5.869 \times 10^{-25}$ | $2.356 \times 10^{-26}$ |
| 17-28 y | $3.320 \times 10^{-26}$ | $7.005 \times 10^{-27}$ |
| 29-36 y | $1.551 \times 10^{-27}$ | $3.149 \times 10^{-28}$ |
| 37-42 y | $1.237 \times 10^{-27}$ | $1.722 \times 10^{-27}$ |
| 43-53 y | $4.990 \times 10^{-28}$ | $1.680 \times 10^{-27}$ |
| 54-66 y | $3.335 \times 10^{-28}$ | $9.691 \times 10^{-28}$ |
| 67-75 y | $2.414 \times 10^{-28}$ | $1.665 \times 10^{-27}$ |
| 76-88 y | $1.126 \times 10^{-28}$ | $3.676 \times 10^{-27}$ |
| 89-101 y | $3.843 \times 10^{-28}$ | $1.599 \times 10^{-26}$ |

**Table S9:** Mann Whitney $p$-values for comparisons between DAST and DGE genes with respect to promoter or gene-body methylation. See Table 1 in the main manuscript for additional information.

Table S10: **Logistic regression results**

| Factor | Estimate | std. error | z value | p-value |
|---|---|---|---|---|
| 1. ZFX | 0.5259 | 0.0935 | 5.6272 | $1.832 \times 10{-}8$ |
| 2. NKX31 | 0.3011 | 0.0803 | 3.7508 | $1.762 \times 10{-}4$ |
| 3. NSD2 | 0.3224 | 0.0870 | 3.7078 | $2.090 \times 10{-}4$ |
| 4. SIX2 | -0.2870 | 0.0787 | -3.6454 | $2.670 \times 10{-}4$ |
| 5. ZEB1 | -0.2719 | 0.0881 | -3.0883 | $2.013 \times 10{-}3$ |
| 6. TF65 | -0.5459 | 0.1799 | -3.0339 | $2.414 \times 10{-}3$ |
| 7. SNAI2 | 0.3114 | 0.1031 | 3.0205 | $2.524 \times 10{-}3$ |
| 8. E2F3 | -0.3707 | 0.1286 | -2.8830 | $3.939 \times 10{-}3$ |
| 9. ATF4 | 0.2267 | 0.0791 | 2.8672 | $4.141 \times 10{-}3$ |
| 10. JARD2 | -0.2475 | 0.0866 | -2.8586 | $4.255 \times 10{-}3$ |
| 11. NKX21 | 0.2666 | 0.0938 | 2.8428 | $4.472 \times 10{-}3$ |
| 12. FOXA2 | 0.2899 | 0.1022 | 2.8361 | $4.567 \times 10{-}3$ |
| 13. DNM3A | 0.3335 | 0.1247 | 2.6752 | $7.468 \times 10{-}3$ |
| 14. KDM5C | -0.2358 | 0.0882 | -2.6722 | $7.535 \times 10{-}3$ |
| 15. FLI1 | -0.3822 | 0.1438 | -2.6571 | $7.882 \times 10{-}3$ |
| 16. ARID2 | 0.2825 | 0.1108 | 2.5483 | 0.0108 |
| 17. SNAI1 | 0.7066 | 0.2864 | 2.4676 | 0.0136 |
| 18. MAD3 | 0.2789 | 0.1139 | 2.4476 | 0.0144 |
| 19. RUNX2 | 0.1958 | 0.0811 | 2.4140 | 0.0158 |
| 20. KAT7 | 0.2375 | 0.0986 | 2.4098 | 0.0160 |
| 21. KDM5A | 0.2634 | 0.1096 | 2.4019 | 0.0163 |
| 22. E2F8 | 0.2572 | 0.1080 | 2.3822 | 0.0172 |
| 23. GATA4 | -0.1998 | 0.0841 | -2.3757 | 0.0175 |
| 24. EMX1 | 0.6204 | 0.2666 | 2.3267 | 0.0200 |
| 25. ID1 | 0.2017 | 0.0868 | 2.3240 | 0.0201 |
| 26. ETV5 | -0.2451 | 0.1090 | -2.2482 | 0.0246 |
| 27. DBP | 0.2781 | 0.1257 | 2.2127 | 0.0269 |
| 28. SIX5 | 0.2468 | 0.1122 | 2.1995 | 0.0278 |
| 29. PPARG | -0.2004 | 0.0927 | -2.1609 | 0.0307 |
| 30. GRHL1 | -0.2385 | 0.1106 | -2.1556 | 0.0311 |
| 31. MAFG | -0.5217 | 0.2423 | -2.1533 | 0.0313 |
| 32. HXD11 | -0.9160 | 0.4318 | -2.1212 | 0.0339 |
| 33. ASCL1 | 0.1685 | 0.0823 | 2.0469 | 0.0407 |
| 34. MYF6 | 0.5928 | 0.2951 | 2.0088 | 0.0446 |
| 35. MYBB | 0.2211 | 0.1106 | 1.9995 | 0.0456 |
| 36. ZBT7B | -0.2155 | 0.1081 | -1.9940 | 0.0461 |
| 37. HNF1B | -0.8304 | 0.4168 | -1.9925 | 0.0463 |
| 38. TBP | -0.3075 | 0.1558 | -1.9739 | 0.0484 |
| 39. ATF1 | 0.2303 | 0.1167 | 1.9735 | 0.0484 |
| 40. HSF1 | -0.1609 | 0.0817 | -1.9686 | 0.0490 |
| 41. EOMES | 0.2598 | 0.1320 | 1.9684 | 0.0490 |
| 42. MXI1 | 0.2313 | 0.1181 | 1.9590 | 0.0501 |
| 43. ELF1 | -0.2980 | 0.1534 | -1.9431 | 0.0520 |
| 44. FOSL1 | -0.1658 | 0.0858 | -1.9315 | 0.0534 |
| 45. UBF1 | 0.1617 | 0.0846 | 1.9115 | 0.0559 |
| 46. KLF15 | 0.2216 | 0.1159 | 1.9114 | 0.0560 |
| 47. CREB1 | 0.2839 | 0.1487 | 1.9092 | 0.0562 |
| 48. E2F6 | -0.2784 | 0.1471 | -1.8924 | 0.0584 |
| 49. PAX5 | 0.2716 | 0.1437 | 1.8903 | 0.0587 |
| 50. HXA4 | 0.1997 | 0.1064 | 1.8762 | 0.0606 |
| 51. MITF | 0.1526 | 0.0814 | 1.8757 | 0.0607 |
| 52. FOSL2 | 0.1709 | 0.0920 | 1.8585 | 0.0631 |

*Continued on next page*

Table S10 – *Continued from previous page*

| Factor | Estimate | std. error | z value | p-value |
|---|---|---|---|---|
| 53. GATA1 | 0.2993 | 0.1645 | 1.8195 | 0.0688 |
| 54. SP4 | 0.1699 | 0.0939 | 1.8094 | 0.0704 |
| 55. PAX2 | 0.3918 | 0.2166 | 1.8088 | 0.0705 |
| 56. ATF6A | -0.2155 | 0.1199 | -1.7972 | 0.0723 |
| 57. TFE3 | -0.4741 | 0.2687 | -1.7646 | 0.0776 |
| 58. MEIS1 | 0.3231 | 0.1848 | 1.7484 | 0.0804 |
| 59. CEBPA | 0.1675 | 0.0959 | 1.7462 | 0.0808 |
| 60. CTCFL | 0.2398 | 0.1378 | 1.7401 | 0.0818 |
| 61. RUNX1 | 0.3203 | 0.1841 | 1.7396 | 0.0819 |
| 62. SPIB | -0.1411 | 0.0811 | -1.7389 | 0.0821 |
| 63. TEAD4 | 0.2005 | 0.1163 | 1.7246 | 0.0846 |
| 64. HXC8 | 0.1914 | 0.1126 | 1.6993 | 0.0893 |
| 65. PRDM1 | 0.1660 | 0.0979 | 1.6957 | 0.0899 |
| 66. SMAD4 | 0.1427 | 0.0850 | 1.6782 | 0.0933 |
| 67. NDF1 | -0.1927 | 0.1153 | -1.6712 | 0.0947 |
| 68. MAFK | 0.1570 | 0.0944 | 1.6629 | 0.0963 |
| 69. ZN250 | -0.2785 | 0.1676 | -1.6617 | 0.0966 |
| 70. BCL3 | 0.1442 | 0.0870 | 1.6573 | 0.0975 |
| 71. KLF9 | 0.1418 | 0.0863 | 1.6422 | 0.1006 |
| 72. CDX2 | 0.1854 | 0.1138 | 1.6296 | 0.1032 |
| 73. TBX3 | 0.6453 | 0.3962 | 1.6285 | 0.1034 |
| 74. NF2L2 | 0.1956 | 0.1210 | 1.6163 | 0.1060 |
| 75. HTF4 | -0.3010 | 0.1873 | -1.6069 | 0.1081 |
| 76. ALX4 | 0.5040 | 0.3145 | 1.6023 | 0.1091 |
| 77. TBPL1 | -0.6871 | 0.4294 | -1.6002 | 0.1095 |
| 78. TFAP4 | -0.1304 | 0.0825 | -1.5814 | 0.1138 |
| 79. DUX4 | 0.1661 | 0.1052 | 1.5784 | 0.1145 |
| 80. KDM5B | -0.2626 | 0.1664 | -1.5779 | 0.1146 |
| 81. ZN236 | -0.7920 | 0.5065 | -1.5637 | 0.1179 |
| 82. P66B | -0.1376 | 0.0890 | -1.5450 | 0.1224 |
| 83. ERG | -0.1387 | 0.0905 | -1.5335 | 0.1252 |
| 84. HXB13 | -0.1681 | 0.1105 | -1.5211 | 0.1282 |
| 85. AP2A | -0.1632 | 0.1082 | -1.5078 | 0.1316 |
| 86. ZIC2 | 0.1724 | 0.1158 | 1.4893 | 0.1364 |
| 87. NF2L3 | -0.7293 | 0.4900 | -1.4882 | 0.1367 |
| 88. ESR2 | -0.1202 | 0.0827 | -1.4531 | 0.1462 |
| 89. ARNT | -0.1439 | 0.1005 | -1.4321 | 0.1521 |
| 90. HEYL | -0.1465 | 0.1038 | -1.4108 | 0.1583 |
| 91. KLF11 | -0.1378 | 0.0978 | -1.4096 | 0.1586 |
| 92. HXA1 | -0.1647 | 0.1176 | -1.3997 | 0.1616 |
| 93. EPAS1 | -0.1641 | 0.1176 | -1.3954 | 0.1629 |
| 94. KLF10 | 0.2615 | 0.1921 | 1.3612 | 0.1734 |
| 95. MSX1 | -0.3144 | 0.2315 | -1.3581 | 0.1744 |
| 96. GLI3 | -0.2132 | 0.1581 | -1.3487 | 0.1774 |
| 97. NFAT5 | 0.1375 | 0.1024 | 1.3422 | 0.1795 |
| 98. BRAC | -0.1089 | 0.0812 | -1.3413 | 0.1798 |
| 99. REST | -0.2058 | 0.1547 | -1.3301 | 0.1835 |
| 100. EGR2 | 0.1258 | 0.0954 | 1.3186 | 0.1873 |
| 101. HNF4A | -0.1486 | 0.1127 | -1.3186 | 0.1873 |
| 102. TSH1 | -0.5508 | 0.4181 | -1.3173 | 0.1877 |
| 103. GATA6 | 0.1491 | 0.1135 | 1.3129 | 0.1892 |
| 104. HXA6 | 0.1318 | 0.1010 | 1.3052 | 0.1918 |
| 105. BMAL1 | -0.1177 | 0.0907 | -1.2984 | 0.1942 |

*Continued on next page*

Table S10 – *Continued from previous page*

| Factor | Estimate | std. error | z value | p-value |
|---|---|---|---|---|
| 106. NR1I2 | -0.3648 | 0.2838 | -1.2854 | 0.1986 |
| 107. ELF3 | -0.2109 | 0.1656 | -1.2735 | 0.2028 |
| 108. HHEX | -0.1709 | 0.1346 | -1.2696 | 0.2042 |
| 109. HIF3A | -0.1177 | 0.0931 | -1.2645 | 0.2061 |
| 110. DLX4 | 0.1803 | 0.1428 | 1.2625 | 0.2068 |
| 111. NCOA3 | -0.3345 | 0.2652 | -1.2614 | 0.2071 |
| 112. FOXA1 | 0.4425 | 0.3511 | 1.2603 | 0.2076 |
| 113. CIC | -0.3441 | 0.2745 | -1.2535 | 0.2100 |
| 114. FOXO1 | -0.1461 | 0.1166 | -1.2527 | 0.2103 |
| 115. GATD1 | -0.1175 | 0.0940 | -1.2508 | 0.2110 |
| 116. ERR1 | 0.1160 | 0.0931 | 1.2465 | 0.2126 |
| 117. EVI1 | -0.1308 | 0.1060 | -1.2345 | 0.2170 |
| 118. DNM3B | 0.1314 | 0.1065 | 1.2341 | 0.2172 |
| 119. RUNX3 | 0.1377 | 0.1125 | 1.2244 | 0.2208 |
| 120. GATA2 | -0.1567 | 0.1282 | -1.2226 | 0.2215 |
| 121. AP2C | -0.1261 | 0.1037 | -1.2164 | 0.2238 |
| 122. P66A | -0.1562 | 0.1287 | -1.2137 | 0.2249 |
| 123. HXA5 | -0.1287 | 0.1061 | -1.2129 | 0.2252 |
| 124. ZBED4 | 0.3275 | 0.2707 | 1.2099 | 0.2263 |
| 125. THA11 | -0.3586 | 0.2997 | -1.1962 | 0.2316 |
| 126. TAL1 | 0.1806 | 0.1510 | 1.1959 | 0.2317 |
| 127. TYY1 | 0.2290 | 0.1917 | 1.1949 | 0.2321 |
| 128. ATF5 | 0.1656 | 0.1388 | 1.1928 | 0.2329 |
| 129. ARI1B | -0.1453 | 0.1226 | -1.1852 | 0.2360 |
| 130. FOXJ2 | 0.2732 | 0.2320 | 1.1778 | 0.2389 |
| 131. LMX1B | 0.4386 | 0.3738 | 1.1734 | 0.2406 |
| 132. IRF2 | 0.1418 | 0.1212 | 1.1697 | 0.2421 |
| 133. JDP2 | -0.2916 | 0.2501 | -1.1660 | 0.2436 |
| 134. KMT2B | 0.1464 | 0.1257 | 1.1649 | 0.2441 |
| 135. MNT | 0.1145 | 0.0989 | 1.1572 | 0.2472 |
| 136. TFDP1 | -0.1241 | 0.1072 | -1.1571 | 0.2472 |
| 137. FOS | 0.1127 | 0.0984 | 1.1451 | 0.2522 |
| 138. GABPA | -0.1689 | 0.1479 | -1.1421 | 0.2534 |
| 139. GMEB2 | 0.1428 | 0.1255 | 1.1380 | 0.2551 |
| 140. CEBPD | 0.1245 | 0.1101 | 1.1314 | 0.2579 |
| 141. ETV7 | -0.1137 | 0.1014 | -1.1211 | 0.2623 |
| 142. USF2 | -0.1155 | 0.1031 | -1.1198 | 0.2628 |
| 143. NFE2 | 0.0943 | 0.0847 | 1.1132 | 0.2656 |
| 144. ELK4 | 0.1138 | 0.1038 | 1.0968 | 0.2727 |
| 145. SRBP2 | -0.3967 | 0.3712 | -1.0684 | 0.2853 |
| 146. MAX | -0.2815 | 0.2638 | -1.0673 | 0.2858 |
| 147. EHF | 0.1104 | 0.1035 | 1.0666 | 0.2861 |
| 148. ZBT7A | -0.1202 | 0.1128 | -1.0658 | 0.2865 |
| 149. LHX2 | 0.1148 | 0.1080 | 1.0630 | 0.2878 |
| 150. MYC | 0.2758 | 0.2610 | 1.0568 | 0.2906 |
| 151. NKX25 | -0.2307 | 0.2194 | -1.0514 | 0.2931 |
| 152. RARG | 0.1321 | 0.1259 | 1.0491 | 0.2941 |
| 153. RELB | 0.0832 | 0.0796 | 1.0450 | 0.2960 |
| 154. GCR | 0.1827 | 0.1758 | 1.0392 | 0.2987 |
| 155. RFX7 | 0.2283 | 0.2226 | 1.0253 | 0.3052 |
| 156. MYB | -0.1457 | 0.1421 | -1.0252 | 0.3053 |
| 157. NDF2 | -0.4349 | 0.4253 | -1.0227 | 0.3065 |
| 158. SALL4 | -0.3440 | 0.3370 | -1.0207 | 0.3074 |

*Continued on next page*

Table S10 – *Continued from previous page*

| Factor | Estimate | std. error | z value | p-value |
|---|---|---|---|---|
| 159. ZFAT | -0.1355 | 0.1330 | -1.0189 | 0.3083 |
| 160. FOXG1 | -0.1674 | 0.1661 | -1.0079 | 0.3135 |
| 161. TEAD1 | -0.0837 | 0.0832 | -1.0049 | 0.3149 |
| 162. HBP1 | -0.2063 | 0.2055 | -1.0041 | 0.3153 |
| 163. LYL1 | -0.0893 | 0.0896 | -0.9962 | 0.3192 |
| 164. RFX1 | 0.1207 | 0.1215 | 0.9936 | 0.3204 |
| 165. NCOR2 | 0.1103 | 0.1113 | 0.9906 | 0.3219 |
| 166. HNF1A | -0.2863 | 0.2911 | -0.9836 | 0.3253 |
| 167. FOSB | 0.3178 | 0.3258 | 0.9753 | 0.3294 |
| 168. CR3L4 | -0.1045 | 0.1073 | -0.9741 | 0.3300 |
| 169. SMRC2 | 0.0865 | 0.0892 | 0.9697 | 0.3322 |
| 170. SMAD5 | -0.2584 | 0.2683 | -0.9629 | 0.3356 |
| 171. MAD1 | 0.1481 | 0.1555 | 0.9523 | 0.3409 |
| 172. NKX61 | 0.2301 | 0.2420 | 0.9509 | 0.3417 |
| 173. RARA | -0.0831 | 0.0882 | -0.9422 | 0.3461 |
| 174. LEF1 | 0.1009 | 0.1073 | 0.9409 | 0.3468 |
| 175. HMBX1 | 0.1362 | 0.1459 | 0.9338 | 0.3504 |
| 176. SIX1 | 0.1309 | 0.1413 | 0.9269 | 0.3540 |
| 177. E2F7 | 0.0904 | 0.0994 | 0.9099 | 0.3629 |
| 178. HXC9 | 0.1120 | 0.1232 | 0.9088 | 0.3635 |
| 179. SFPQ | -0.0860 | 0.0947 | -0.9078 | 0.3640 |
| 180. NKX22 | 0.0973 | 0.1093 | 0.8900 | 0.3735 |
| 181. SOX10 | 0.1288 | 0.1464 | 0.8797 | 0.3790 |
| 182. ZN639 | 0.2339 | 0.2669 | 0.8764 | 0.3808 |
| 183. ZN263 | 0.0918 | 0.1048 | 0.8757 | 0.3812 |
| 184. YBOX3 | 0.4226 | 0.4851 | 0.8713 | 0.3836 |
| 185. MEIS2 | -0.1595 | 0.1842 | -0.8661 | 0.3864 |
| 186. HNF4G | -0.0740 | 0.0855 | -0.8646 | 0.3873 |
| 187. BC11A | -0.0812 | 0.0945 | -0.8593 | 0.3902 |
| 188. ZN335 | 0.2714 | 0.3178 | 0.8541 | 0.3931 |
| 189. MAFF | 0.0974 | 0.1145 | 0.8506 | 0.3950 |
| 190. PITX1 | 0.1984 | 0.2334 | 0.8501 | 0.3953 |
| 191. HXC11 | -0.3675 | 0.4397 | -0.8360 | 0.4032 |
| 192. GLIS1 | -0.0776 | 0.0932 | -0.8332 | 0.4047 |
| 193. SOX9 | 0.1248 | 0.1516 | 0.8231 | 0.4104 |
| 194. NFYC | 0.0999 | 0.1215 | 0.8222 | 0.4110 |
| 195. SMAD3 | -0.0787 | 0.0958 | -0.8211 | 0.4116 |
| 196. NRF1 | 0.1013 | 0.1254 | 0.8078 | 0.4192 |
| 197. STAT1 | -0.0807 | 0.1002 | -0.8057 | 0.4204 |
| 198. RFX5 | -0.0763 | 0.0948 | -0.8052 | 0.4207 |
| 199. HXB6 | -0.2436 | 0.3029 | -0.8043 | 0.4212 |
| 200. PO5F1 | 0.0936 | 0.1176 | 0.7952 | 0.4265 |
| 201. THAP1 | 0.0806 | 0.1019 | 0.7914 | 0.4287 |
| 202. E2F1 | -0.1139 | 0.1442 | -0.7897 | 0.4297 |
| 203. ERF | 0.1608 | 0.2061 | 0.7803 | 0.4352 |
| 204. MAZ | 0.1966 | 0.2529 | 0.7776 | 0.4368 |
| 205. UBIP1 | 0.0931 | 0.1199 | 0.7766 | 0.4374 |
| 206. JUND | 0.1133 | 0.1475 | 0.7683 | 0.4423 |
| 207. ELF5 | -0.1244 | 0.1632 | -0.7623 | 0.4459 |
| 208. GLIS3 | -0.1666 | 0.2197 | -0.7583 | 0.4482 |
| 209. ISL1 | -0.1160 | 0.1545 | -0.7507 | 0.4529 |
| 210. P73 | -0.0603 | 0.0804 | -0.7494 | 0.4536 |
| 211. FOXM1 | -0.0972 | 0.1299 | -0.7488 | 0.4540 |

*Continued on next page*

Table S10 – *Continued from previous page*

| Factor | Estimate | std. error | z value | p-value |
|---|---|---|---|---|
| 212. FOXK1 | -0.0724 | 0.0973 | -0.7444 | 0.4566 |
| 213. IRF3 | -0.1177 | 0.1587 | -0.7414 | 0.4585 |
| 214. TEAD2 | -0.0865 | 0.1169 | -0.7400 | 0.4593 |
| 215. CR3L2 | -0.2512 | 0.3412 | -0.7361 | 0.4617 |
| 216. HXA10 | 0.1652 | 0.2249 | 0.7346 | 0.4626 |
| 217. RORA | -0.1771 | 0.2422 | -0.7312 | 0.4646 |
| 218. BACH1 | -0.0804 | 0.1103 | -0.7294 | 0.4658 |
| 219. ATF2 | 0.0740 | 0.1031 | 0.7177 | 0.4729 |
| 220. ARNT2 | 0.0922 | 0.1292 | 0.7138 | 0.4753 |
| 221. ZNF83 | -0.0667 | 0.0937 | -0.7118 | 0.4766 |
| 222. ESR1 | -0.1446 | 0.2037 | -0.7098 | 0.4778 |
| 223. ZN266 | 0.0794 | 0.1122 | 0.7076 | 0.4792 |
| 224. PATZ1 | -0.1325 | 0.1876 | -0.7063 | 0.4800 |
| 225. MYCN | 0.0859 | 0.1220 | 0.7046 | 0.4811 |
| 226. SOX4 | 0.1061 | 0.1512 | 0.7017 | 0.4829 |
| 227. VEZF1 | -0.0703 | 0.1012 | -0.6940 | 0.4877 |
| 228. E2F4 | 0.0780 | 0.1124 | 0.6939 | 0.4878 |
| 229. MEF2B | -0.0685 | 0.0989 | -0.6920 | 0.4889 |
| 230. FOXO3 | -0.1414 | 0.2049 | -0.6902 | 0.4901 |
| 231. BCL6 | 0.0926 | 0.1344 | 0.6891 | 0.4908 |
| 232. TAF1 | -0.1096 | 0.1603 | -0.6837 | 0.4942 |
| 233. ARI1A | -0.0720 | 0.1054 | -0.6830 | 0.4946 |
| 234. ETS1 | 0.1042 | 0.1536 | 0.6784 | 0.4975 |
| 235. KLF1 | 0.0757 | 0.1120 | 0.6756 | 0.4993 |
| 236. PBX2 | -0.0890 | 0.1320 | -0.6740 | 0.5003 |
| 237. NR1H3 | -0.0726 | 0.1078 | -0.6739 | 0.5004 |
| 238. DLX6 | -0.1172 | 0.1770 | -0.6619 | 0.5080 |
| 239. STAT2 | 0.0694 | 0.1055 | 0.6581 | 0.5105 |
| 240. AEBP2 | -0.0861 | 0.1336 | -0.6443 | 0.5194 |
| 241. PO2F2 | 0.0709 | 0.1105 | 0.6420 | 0.5209 |
| 242. TOX4 | -0.1873 | 0.2930 | -0.6393 | 0.5226 |
| 243. PBX1 | 0.1015 | 0.1590 | 0.6385 | 0.5232 |
| 244. GFI1B | -0.0797 | 0.1267 | -0.6293 | 0.5292 |
| 245. SOX2 | -0.0705 | 0.1126 | -0.6262 | 0.5312 |
| 246. NR1H2 | -0.0578 | 0.0934 | -0.6191 | 0.5358 |
| 247. MZF1 | -0.0831 | 0.1343 | -0.6188 | 0.5361 |
| 248. KLF3 | -0.0710 | 0.1163 | -0.6104 | 0.5416 |
| 249. Z280D | -0.0720 | 0.1184 | -0.6079 | 0.5433 |
| 250. HXB4 | 0.0983 | 0.1626 | 0.6048 | 0.5453 |
| 251. HES1 | -0.0705 | 0.1166 | -0.6045 | 0.5455 |
| 252. BHE40 | 0.0566 | 0.0947 | 0.5973 | 0.5503 |
| 253. COT2 | -0.0512 | 0.0858 | -0.5967 | 0.5507 |
| 254. ZEP1 | -0.1529 | 0.2576 | -0.5938 | 0.5526 |
| 255. CDC5L | 0.2420 | 0.4092 | 0.5914 | 0.5542 |
| 256. STAT4 | -0.0684 | 0.1159 | -0.5899 | 0.5552 |
| 257. ID2 | -0.1274 | 0.2168 | -0.5874 | 0.5569 |
| 258. MYNN | -0.0849 | 0.1458 | -0.5821 | 0.5605 |
| 259. DLX1 | -0.0640 | 0.1100 | -0.5818 | 0.5607 |
| 260. TF7L2 | -0.0517 | 0.0890 | -0.5807 | 0.5614 |
| 261. HXA13 | 0.3466 | 0.6045 | 0.5733 | 0.5665 |
| 262. HXA9 | 0.2274 | 0.4077 | 0.5577 | 0.5771 |
| 263. FOXP1 | -0.0604 | 0.1096 | -0.5507 | 0.5818 |
| 264. PROX1 | -0.0525 | 0.0957 | -0.5485 | 0.5834 |

*Continued on next page*

Table S10 – *Continued from previous page*

| Factor | Estimate | std. error | z value | p-value |
|---|---|---|---|---|
| 265. P63 | 0.0597 | 0.1096 | 0.5449 | 0.5858 |
| 266. PLAG1 | 0.1733 | 0.3221 | 0.5379 | 0.5907 |
| 267. RXRA | -0.0746 | 0.1394 | -0.5348 | 0.5928 |
| 268. EGR1 | 0.0620 | 0.1165 | 0.5318 | 0.5949 |
| 269. NCOR1 | 0.0539 | 0.1032 | 0.5228 | 0.6011 |
| 270. GMEB1 | 0.0619 | 0.1253 | 0.4943 | 0.6211 |
| 271. MEF2D | 0.2120 | 0.4311 | 0.4917 | 0.6229 |
| 272. GATA3 | 0.0609 | 0.1261 | 0.4832 | 0.6290 |
| 273. ZBT17 | 0.0404 | 0.0842 | 0.4798 | 0.6313 |
| 274. OTX2 | -0.0881 | 0.1842 | -0.4785 | 0.6323 |
| 275. ATRX | 0.0457 | 0.0962 | 0.4755 | 0.6345 |
| 276. SP2 | 0.0476 | 0.1005 | 0.4739 | 0.6356 |
| 277. MTA3 | 0.0873 | 0.1872 | 0.4664 | 0.6409 |
| 278. SRBP1 | 0.0481 | 0.1054 | 0.4566 | 0.6480 |
| 279. BARX2 | 0.0554 | 0.1219 | 0.4544 | 0.6496 |
| 280. SMAD1 | -0.0406 | 0.0895 | -0.4531 | 0.6505 |
| 281. STAT6 | -0.0446 | 0.1013 | -0.4410 | 0.6592 |
| 282. TFCP2 | -0.1726 | 0.3965 | -0.4354 | 0.6632 |
| 283. PRD14 | -0.0359 | 0.0826 | -0.4346 | 0.6638 |
| 284. MYBA | 0.0725 | 0.1671 | 0.4341 | 0.6642 |
| 285. IRF1 | -0.0528 | 0.1223 | -0.4315 | 0.6661 |
| 286. CEBPB | -0.1127 | 0.2650 | -0.4252 | 0.6707 |
| 287. ETV1 | -0.0430 | 0.1011 | -0.4249 | 0.6709 |
| 288. ZN281 | 0.0418 | 0.0986 | 0.4243 | 0.6713 |
| 289. ITF2 | -0.0377 | 0.0893 | -0.4220 | 0.6730 |
| 290. ID3 | 0.0452 | 0.1071 | 0.4219 | 0.6731 |
| 291. SRF | 0.0419 | 0.0998 | 0.4199 | 0.6745 |
| 292. CASZ1 | 0.0853 | 0.2037 | 0.4186 | 0.6755 |
| 293. ZFHX3 | 0.0457 | 0.1103 | 0.4139 | 0.6790 |
| 294. PPARA | 0.1205 | 0.2941 | 0.4099 | 0.6819 |
| 295. IKZF1 | -0.0343 | 0.0843 | -0.4067 | 0.6842 |
| 296. MAF | -0.0550 | 0.1364 | -0.4030 | 0.6869 |
| 297. SOX3 | 0.1295 | 0.3257 | 0.3975 | 0.6910 |
| 298. KLF5 | 0.0513 | 0.1304 | 0.3934 | 0.6940 |
| 299. PRGR | -0.0661 | 0.1693 | -0.3903 | 0.6963 |
| 300. PO2F1 | -0.0626 | 0.1603 | -0.3903 | 0.6963 |
| 301. HES4 | 0.0614 | 0.1575 | 0.3897 | 0.6968 |
| 302. HIF1A | 0.0333 | 0.0875 | 0.3804 | 0.7036 |
| 303. XBP1 | 0.0383 | 0.1020 | 0.3754 | 0.7073 |
| 304. RFX2 | 0.0376 | 0.1007 | 0.3733 | 0.7089 |
| 305. SMRC1 | 0.0384 | 0.1037 | 0.3708 | 0.7108 |
| 306. FOXP3 | 0.0349 | 0.0959 | 0.3640 | 0.7158 |
| 307. ASCL2 | 0.0400 | 0.1099 | 0.3637 | 0.7161 |
| 308. TWST1 | -0.0330 | 0.0910 | -0.3628 | 0.7167 |
| 309. SUH | 0.0408 | 0.1145 | 0.3559 | 0.7219 |
| 310. NR4A1 | 0.0362 | 0.1021 | 0.3550 | 0.7226 |
| 311. ETV4 | 0.0507 | 0.1434 | 0.3535 | 0.7237 |
| 312. MEF2C | 0.0366 | 0.1046 | 0.3504 | 0.7260 |
| 313. SPI1 | 0.0844 | 0.2424 | 0.3481 | 0.7278 |
| 314. CLOCK | 0.0405 | 0.1166 | 0.3473 | 0.7284 |
| 315. SP1 | 0.0664 | 0.1915 | 0.3471 | 0.7286 |
| 316. CREM | 0.0578 | 0.1677 | 0.3445 | 0.7305 |
| 317. ATOH1 | -0.0651 | 0.1913 | -0.3404 | 0.7336 |

*Continued on next page*

Table S10 – *Continued from previous page*

| Factor | Estimate | std. error | z value | p-value |
|---|---|---|---|---|
| 318. SOX15 | -0.1620 | 0.4912 | -0.3297 | 0.7416 |
| 319. KDM5D | -0.0297 | 0.0903 | -0.3292 | 0.7420 |
| 320. ZN407 | 0.0484 | 0.1472 | 0.3289 | 0.7422 |
| 321. CTCF | 0.0870 | 0.2664 | 0.3265 | 0.7441 |
| 322. ZN217 | -0.0334 | 0.1027 | -0.3252 | 0.7451 |
| 323. AHR | 0.0764 | 0.2358 | 0.3241 | 0.7459 |
| 324. SP3 | -0.1256 | 0.3910 | -0.3212 | 0.7480 |
| 325. KLF6 | -0.0570 | 0.1782 | -0.3201 | 0.7489 |
| 326. MYOD1 | -0.0281 | 0.0878 | -0.3197 | 0.7492 |
| 327. NANOG | 0.0266 | 0.0840 | 0.3167 | 0.7515 |
| 328. COE1 | 0.0340 | 0.1082 | 0.3140 | 0.7535 |
| 329. E2F2 | -0.0354 | 0.1144 | -0.3094 | 0.7570 |
| 330. ZNF84 | 0.0313 | 0.1070 | 0.2929 | 0.7696 |
| 331. BARX1 | -0.0365 | 0.1332 | -0.2742 | 0.7840 |
| 332. SMAD2 | -0.0239 | 0.0873 | -0.2739 | 0.7842 |
| 333. ZN143 | 0.0278 | 0.1039 | 0.2679 | 0.7888 |
| 334. NR2C2 | 0.0227 | 0.0862 | 0.2628 | 0.7927 |
| 335. PBX3 | -0.0252 | 0.0963 | -0.2621 | 0.7933 |
| 336. BARH1 | 0.0371 | 0.1432 | 0.2593 | 0.7954 |
| 337. NFAC3 | -0.0826 | 0.3192 | -0.2588 | 0.7958 |
| 338. IRX2 | -0.0418 | 0.1627 | -0.2567 | 0.7974 |
| 339. BATF | 0.0225 | 0.0878 | 0.2561 | 0.7979 |
| 340. KMT2A | 0.0228 | 0.0891 | 0.2554 | 0.7984 |
| 341. FOXO4 | -0.0826 | 0.3241 | -0.2548 | 0.7989 |
| 342. ATF3 | -0.0291 | 0.1190 | -0.2450 | 0.8065 |
| 343. NR5A2 | 0.0788 | 0.3289 | 0.2396 | 0.8106 |
| 344. KLF4 | 0.0197 | 0.0825 | 0.2382 | 0.8117 |
| 345. KAT5 | -0.0300 | 0.1270 | -0.2361 | 0.8133 |
| 346. OVOL2 | -0.0402 | 0.1703 | -0.2361 | 0.8133 |
| 347. KAISO | 0.0243 | 0.1031 | 0.2355 | 0.8138 |
| 348. FOXH1 | 0.0190 | 0.0808 | 0.2353 | 0.8139 |
| 349. MIXL1 | -0.0448 | 0.1933 | -0.2317 | 0.8168 |
| 350. FEV | 0.0235 | 0.1047 | 0.2245 | 0.8224 |
| 351. TCF7 | -0.0438 | 0.2025 | -0.2164 | 0.8286 |
| 352. FOXC1 | 0.0555 | 0.2627 | 0.2114 | 0.8325 |
| 353. DLX2 | 0.0409 | 0.1972 | 0.2073 | 0.8358 |
| 354. CEBPG | 0.0376 | 0.1918 | 0.1962 | 0.8444 |
| 355. P53 | 0.0271 | 0.1433 | 0.1888 | 0.8502 |
| 356. HXC6 | -0.0258 | 0.1384 | -0.1860 | 0.8524 |
| 357. NFAC1 | 0.0196 | 0.1070 | 0.1834 | 0.8545 |
| 358. FOXP2 | -0.0195 | 0.1066 | -0.1833 | 0.8546 |
| 359. USF1 | 0.0258 | 0.1467 | 0.1759 | 0.8604 |
| 360. NFIB | -0.0662 | 0.3975 | -0.1666 | 0.8677 |
| 361. ZNF92 | 0.0139 | 0.0840 | 0.1660 | 0.8682 |
| 362. NCOA1 | 0.0146 | 0.0896 | 0.1634 | 0.8702 |
| 363. NFIC | -0.0138 | 0.0878 | -0.1568 | 0.8754 |
| 364. ELF2 | -0.0203 | 0.1310 | -0.1549 | 0.8769 |
| 365. DMRT1 | -0.0256 | 0.1672 | -0.1533 | 0.8782 |
| 366. LHX4 | -0.0473 | 0.3213 | -0.1472 | 0.8830 |
| 367. STA5A | -0.0130 | 0.0957 | -0.1363 | 0.8916 |
| 368. ZBT44 | -0.0578 | 0.4346 | -0.1329 | 0.8943 |
| 369. TF7L1 | 0.0116 | 0.0878 | 0.1326 | 0.8945 |
| 370. TBX21 | -0.0117 | 0.0894 | -0.1309 | 0.8958 |

*Continued on next page*

Table S10 – *Continued from previous page*

| Factor | Estimate | std. error | z value | p-value |
|---|---|---|---|---|
| 371. HXC13 | 0.0524 | 0.4203 | 0.1248 | 0.9007 |
| 372. NKX23 | 0.0159 | 0.1274 | 0.1246 | 0.9008 |
| 373. TFE2 | -0.0144 | 0.1161 | -0.1240 | 0.9013 |
| 374. PDX1 | -0.0103 | 0.0934 | -0.1102 | 0.9122 |
| 375. SATB1 | -0.0451 | 0.4139 | -0.1090 | 0.9132 |
| 376. STF1 | 0.0487 | 0.4608 | 0.1057 | 0.9158 |
| 377. HINFP | -0.0118 | 0.1139 | -0.1035 | 0.9175 |
| 378. NFKB1 | 0.0077 | 0.0790 | 0.0972 | 0.9226 |
| 379. ZBT16 | -0.0207 | 0.2135 | -0.0971 | 0.9226 |
| 380. BACH2 | 0.0083 | 0.0915 | 0.0906 | 0.9278 |
| 381. IRF4 | 0.0099 | 0.1108 | 0.0897 | 0.9285 |
| 382. NFKB2 | -0.0075 | 0.0836 | -0.0893 | 0.9288 |
| 383. GRHL2 | -0.0081 | 0.0936 | -0.0861 | 0.9314 |
| 384. JUNB | 0.0079 | 0.0920 | 0.0861 | 0.9314 |
| 385. JUN | -0.0116 | 0.1359 | -0.0853 | 0.9321 |
| 386. HSF2 | 0.0223 | 0.2739 | 0.0815 | 0.9350 |
| 387. MEF2A | 0.0067 | 0.0894 | 0.0746 | 0.9405 |
| 388. HXA2 | -0.0069 | 0.0979 | -0.0708 | 0.9436 |
| 389. ANDR | 0.0217 | 0.3068 | 0.0707 | 0.9437 |
| 390. ZBTB2 | 0.0142 | 0.2016 | 0.0704 | 0.9439 |
| 391. STAT3 | 0.0086 | 0.1349 | 0.0637 | 0.9492 |
| 392. CUX1 | -0.0123 | 0.2221 | -0.0555 | 0.9557 |
| 393. FOXD2 | 0.0050 | 0.1011 | 0.0495 | 0.9605 |
| 394. GLI1 | -0.0086 | 0.2033 | -0.0422 | 0.9663 |
| 395. GLI2 | -0.0048 | 0.1132 | -0.0422 | 0.9664 |
| 396. ZN274 | -0.0058 | 0.1487 | -0.0393 | 0.9686 |
| 397. NFYA | 0.0022 | 0.1085 | 0.0200 | 0.9840 |
| 398. COT1 | 0.0023 | 0.1700 | 0.0134 | 0.9893 |
| 399. VDR | 0.0012 | 0.1162 | 0.0105 | 0.9916 |
| 400. NFYB | 0.0009 | 0.1067 | 0.0089 | 0.9929 |
| 401. CEBPE | -0.0015 | 0.4016 | -0.0038 | 0.9970 |

### References


1. Grossmann, S., Bauer, S., Robinson, P. N., and Vingron, M. Improved detection of overrepresentation of gene-ontology annotations with parent child analysis. *Bioinformatics (Oxford, England)* **23**, 3024–3031, November (2007).

2. Bauer, S., Grossmann, S., Vingron, M., and Robinson, P. N. Ontologizer 2.0–a multifunctional tool for GO term enrichment analysis and data exploration. *Bioinformatics (Oxford, England)* **24**, 1650–1651, July (2008).
